# Systems analysis shows a role of cytophilic antibodies in shaping innate tolerance to malaria

**DOI:** 10.1101/2021.09.02.458668

**Authors:** Maximilian Julius Lautenbach, Victor Yman, Nadir Kadri, David Fernando Plaza, Sina Angenendt, Klara Sondén, Anna Färnert, Christopher Sundling

## Abstract

The mechanism of acquisition and maintenance of natural immunity against *Plasmodium falciparum* malaria remains unclear. Although, clinical immunity develops over time with repeated malaria episodes, disease tolerance is more rapidly acquired compared to protective immunity. It remains unclear, how pre-existing immune responses impacts the mechanism responsible for disease tolerance. Here, we investigated a cohort of returning travelers treated for acute symptomatic *P. falciparum* malaria, either infected for the first time, or with a previous history of malaria. Through repeated sampling over one year in a malaria free setting, we were able to study the acute and longitudinal effects of the infection. We combined comprehensive immune cell and plasma protein profiling with integrated and data driven analysis, describing the immune landscape from acute disease to one year after infection. We identified a strong association between pro-inflammatory signatures and γδ T cell expansion. The association was significantly impacted by previous exposure to malaria, resulting in a dampened pro-inflammatory response, which translated to reduced Vδ2^+^ γδ T cell expansion compared to primary infected individuals. The dampened inflammatory signal was associated with early expansion of FcγRIII+ monocytes and parasite-specific antibodies of IgG1 and IgG3 isotypes.

Our data suggest that the interplay of FcγRIII+ monocytes and a cytophilic parasite-specific IgG during the early blood stage infection lead to lower parasitemia and a dampened pro-inflammatory response with reduced γδ T cell expansion. This enhanced control and reduced inflammation points to a potential mechanism on how tolerance is established following repeated malaria exposure.

**One Sentence Summary:** A systems immunology analysis on natural malaria sheds light on disease tolerance mechanism associated with gamma delta T cell expansion

## INTRODUCTION

Malaria remains a global burden with an estimated 229 million malaria cases leading to approximately 409 000 deaths in 2019 *(1)*. Compared to other pathogens, immunity to malaria is slow to develop *(2)*. In malaria endemic areas, partial immunity is acquired over time after repeated infections. Clinical symptoms are reduced while low grade subclinical infections remain and allow for continued parasite transmission *(2, 3)*. However, protection from severe forms of malaria, despite high levels of parasitemia, seem to develop already after a few infections *(4)*. A recent study by Nahrendorf and colleagues suggests a tolerance mechanism where better control of host-damaging factors that are part of the natural immune response to infection are an underlying cause for the reduced severity *(5)*. Currently no study has investigated how long tolerance persists and how it is maintained in response to malaria.

Clinical symptoms are due to the blood stage of the infection, where merozoites invade erythrocytes that then sequester, followed by parasite multiplication before bursting out to infect new erythrocytes *(6)*. The clinical manifestations are partly due to the strong innate pro-inflammatory response, during the parasite blood stage/to the *P. falciparum* blood stage *(7)*. Much of our current understanding of the immune responses to *P. falciparum* infection is based on hypothesis-driven research, where selected cell subsets, inflammatory markers, or clinical features are investigated at a time *(8, 9)*. Improved technology and bioinformatic tools enable the analysis of high-dimensional parameters in limited sample material, giving us the opportunity to study complex immune responses at a systems level to provide new insights into complex immunological networks that represent the immune response during infection *(10)*. Systems level approaches to study febrile/non-febrile children in malaria endemic countries *(11)* as well as the comparison of malaria naïve and Africans in controlled human malaria infection (CHMI) studies *(12)* have generated valuable findings and new hypotheses. While studies in endemic populations allow investigation of acute naturally acquired malaria and persistent (largely asymptomatic) infections in the context of previous exposure. Despite CHMI studies enable more control over prior parasite exposure, infectious dose, and follow-up after infection, although only in the context of very low level parasitemia and often before symptoms having appeared. Although compensating for some limitations, CHMI cannot fully mirror a natural infection, due to early treatment not allowing observation of potential effects derived from a strong natural symptomatic infection. This leaves a knowledge gap on how natural infection affects the immune response in the absence of potential parasite re-exposure. This gap can be filled by investigating the immune response after naturally acquired malaria in individuals leaving the endemic area, seeking healthcare in a setting without risk of re-exposure.

Here, we study a prospective cohort of returning travelers treated for acute *P. falciparum* malaria at the Karolinska University Hospital in Sweden, followed over one year after infection. Using a systems immunology approach and data-driven analysis, we combined plasma protein and cell profiling to study dynamic changes of the immune response over time in these patients. Based on our results, we propose a model where antibody-derived memory modulates the pro-inflammatory cytokine response which in turn impact γδ T cell expansion and improve disease tolerance.

## RESULTS

### Subhead 1: Longitudinal profiling of peripheral blood – Immune dynamics after natural infection in a cohort of returning travelers

We aimed to comprehensively profile the immune response dynamics longitudinally after natural infection with *P. falciparum* malaria, to appreciate the immunological changes occurring during the acute disease and up to one year after diagnosis. We included 53 returning travelers, repeatedly sampled in a non-endemic setting over one year after hospital admission *(13–16)* (Fig. 1A).

**Fig. 1.**
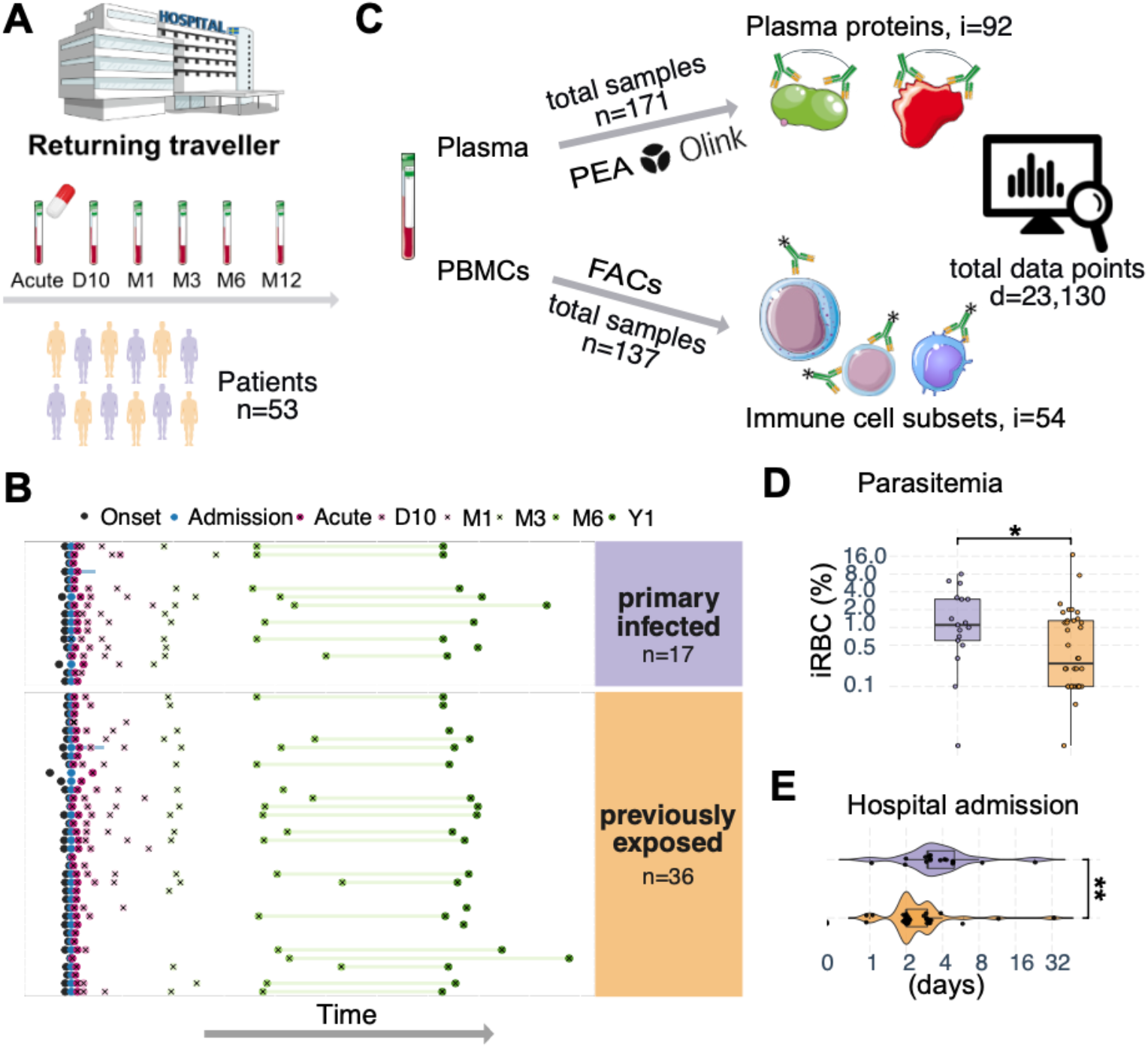
Systems level profiling of peripheral blood from individuals of a prospective cohort of returning travelers with P. falciparum malaria. **(A)** Overview of prospective longitudinal malaria cohort of returning travelers, **(B)** temporal sample information symptom onset (black), admission to Karolinska University Hospital (blue), six sampling time points (x) for immune profiling (acute, after 10 days, 1 month, 3 months, 6 months, 1 year) and convalescence average (green lines). Patients with primary infection (n = 17) are colored in purple, and patients with previous malaria exposure (n = 36) are colored in orange. **(C)** A total of 53 returning travelers were longitudinally profiled for plasma proteins using Olink Proteomics PEA platform (n = 171 samples, i = 92 proteins) and their PBMC immune cells (n = 137 samples, i = 54 subsets), generating 23,130 unique data points. Comparison of **(D)** parasitemia at diagnosis (percentage of infected red blood cells) and **(E)** length of hospital admission (days) for primary infected and previously exposed individuals. Statistical differences between groups were assessed using the non-parametric Wilcoxon-test. *p < 0.05, **p < 0.01.

To profile the immune response in the cohort over time, we performed broad FACS-based immunophenotyping using a 17-marker panel and gated for subsets of monocytes (CD14^+^), T cells (CD3^+^CD56^−^), and NK cell (CD3^−^CD56^+^) (Fig. S1A). We also included data from B cell subsets previously phenotyped from the same donors *(16)* (Fig. S1B). Plasma protein analysis was performed using the Target 96-plex Inflammation panel from Olink Proteomics *(17)* targeting 92 immune response-related proteins. In total, we profiled 182 samples from 53 subjects, creating more than 23,000 data points (Fig. 1B).

The cohort consisted of individuals infected with *P. falciparum* malaria for the first time (primary infected, n = 17) and those infected before, having grown up in malaria endemic areas and reported previous malaria (previously exposed, n = 36) (Fig. 1C). Comparing individuals with primary infection versus previous exposure enabled us to investigate potential memory effect on the immune response. The individuals with previous exposure moved from malaria endemic areas to non-endemic Sweden on average 11.5 years before now experiencing acute malaria after visiting an endemic area (median = 11.5, range 0-46 years, Tab. S1).

Although all individuals sought healthcare, primary infected patients had significantly higher levels of parasitemia at hospital admission (median 1.10 vs 0.25 % infected red blood cells, p = 0.038, Fig1D), more signs of severe malaria (Tab. S1) and were on average admitted to hospital care significantly longer than previously exposed patients (median 3 vs 2 days, p = 0.002, Fig1E, Tab. S1).

### Subhead 2: Integrated analysis of immune response on time axis after symptom onset

To explore the immune response dynamics after natural *P. falciparum* malaria on a systems level and in a data driven manner, we used multi-omics factor analysis (MOFA). The unsupervised nature of MOFA allows the model to capture both biological and technical variability in the low-dimensional factors space *(18, 19)*. Here we used MEFISTO, a recent extension of MOFA which allows for multi-omics integration of data while controlling for time-dependent variance in repeated samples *(20)*. MOFA can disentangle the sources of heterogeneity in diverse data types and accept missing data points using matrix factorization (Fig. S2A). Using MEFISTO, the dataset was described by two latent factors. These factors explained 37% to 60% and 58% to 72 % of the immune cell subset and plasma protein dynamics, respectively (Fig. 2A). Factor1 was primarily associated with time-dependent changes in the immune response to the infection (Fig. 2B and S2B). Positive factor values were associated with the acute phase and negative factor values were associated with the time after treatment and transition towards convalescence (Fig. 2B and S2C). The positive factor values were driven by increased levels of IL10, pro-inflammatory cytokines such as IFN-gamma, IL-6, TNF, CCL3, IL8, CDCP1, and chemokines such as CXCL9, CXCL10, CXCL11. For immune cells, CD4^+^ and CD8^+^ T cells expressing activation markers CD38 and HLA-DR, intermediate monocytes (CD14^+^CD16^+^) and plasmablasts (CD19^+^CD20^lo^CD38^hi^CD27^hi^) were associated with Factor1 (Fig. 2C). The negative factor values were associated with the transition phase towards convalescence. Here, several subsets of NK cells expressing CD57, CD8 T effector memory cells, and γδ T cells were variables driving Factor1 for the cellular immune response after treatment (Fig. 2C). We confirmed the accuracy of the model parameters by comparing measured data between the acute and convalescent samples (internal control; Fig. S2C-F) and to healthy control samples (external control; Fig. S3A-D).

**Fig. 2.**
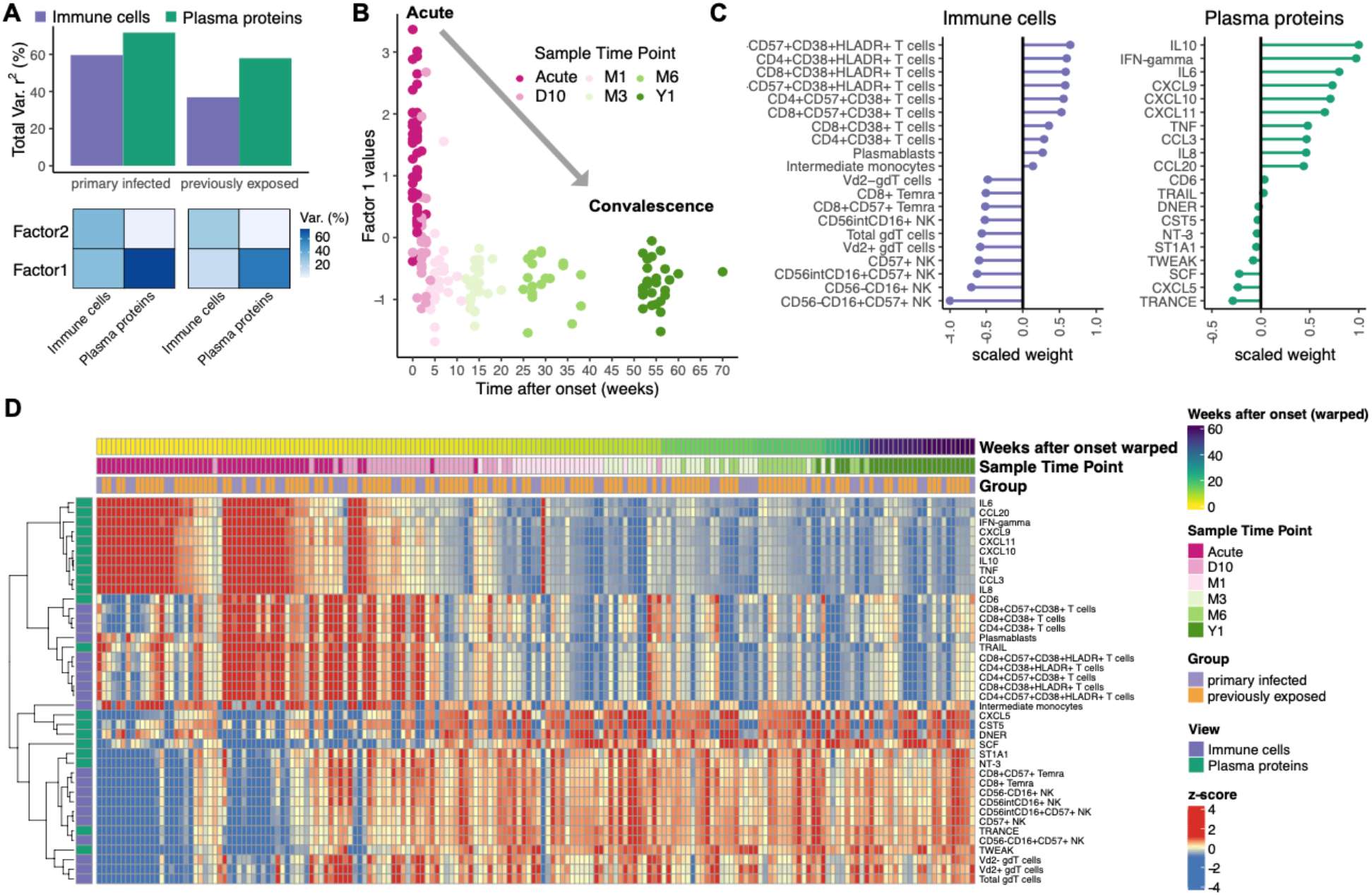
Integrated analysis of immune response dynamics for one year after malaria. Integrated multi omics factor analysis method for the Functional Integration of Spatial and Temporal Omics data (MEFISTO) model was utilized to integrate data modalities, plasma proteins and manually gated cell subsets, respectively, with their temporal covariate of time after symptom onset. (**A**) Variance explained by the MEFISTO model. Differences due to previous parasite exposure were assigned to groups in order to model group-specific time-dependent immune response dynamics. (**B**) Plotting Factor1 value against the time after symptom onset (weeks) highlights the captured variance in Factor1 with positive factor values associated with acute phase samples and negative factor values associated with non-acute phase samples. (**C**) Top10 features of Factor1 for each of the immune system views based on negative and positive scaled feature weights - immune cells (purple) and plasma proteins (green). (**D)** The heatmaps visualize immune response dynamics for Factor1 driving features. The characteristic immune variables for each data modality, aligned to time after symptom onset, reveals contrasting patterns characterizing the transition from acute phase towards convalescence. Missing data points have been imputed based on model Factor1. Cell counts and relative protein levels are shown as rows, and each column represents an individual patient. Rows were clustered using Euclidean distance and column were arranged according to MEFISTO model group-aligned time after onset. Cell counts and protein level values were converted to z-scores.

We then used the integrated MEFISTO model to impute missing data points (plasma proteins data modality, n = 11; immune cell modality, n = 64; Fig. S2A) to generate a heatmap of the integrated immune landscape, visualizing the longitudinal immune system dynamics after disease. The landscape dynamics, show a rapidly contracting pro-inflammatory response after treatment with a temporary increase in primarily activated T cell subsets followed by a transition into a more long-term post-infection response (Fig. 2D and Fig. S3).

In summary, our integrated analysis of immune cell subsets and plasma protein data after symptom onset allows us to draw a data driven descriptive immune landscape, describing the transition from the clinical acute phase towards convalescence over one year.

### Subhead 3: Relationship between acute cytokine milieu and cellular response

Cytokines and chemokines are known to orchestrate an immunological response to drive recruitment, activation, and subsequently proliferation of immune cell populations. Hence, we sought to analyze possible associations between levels of the acute phase responses and immune cells subsets, characterizing the transition from acute towards convalescence.

Focusing on the top immune variables determined by the integrated model, we correlated acute samples with all remaining sample time points using spearman rank correlation (Fig. 3A). We discovered strong and significant correlations (|rho| > 0.7, after adjusted FDR < 0.05) between acute phase cytokines and immune cell subsets at all sample time points after treatment (D10, M3, M6 and Y1) (Fig. 3B).

**Fig. 3.**
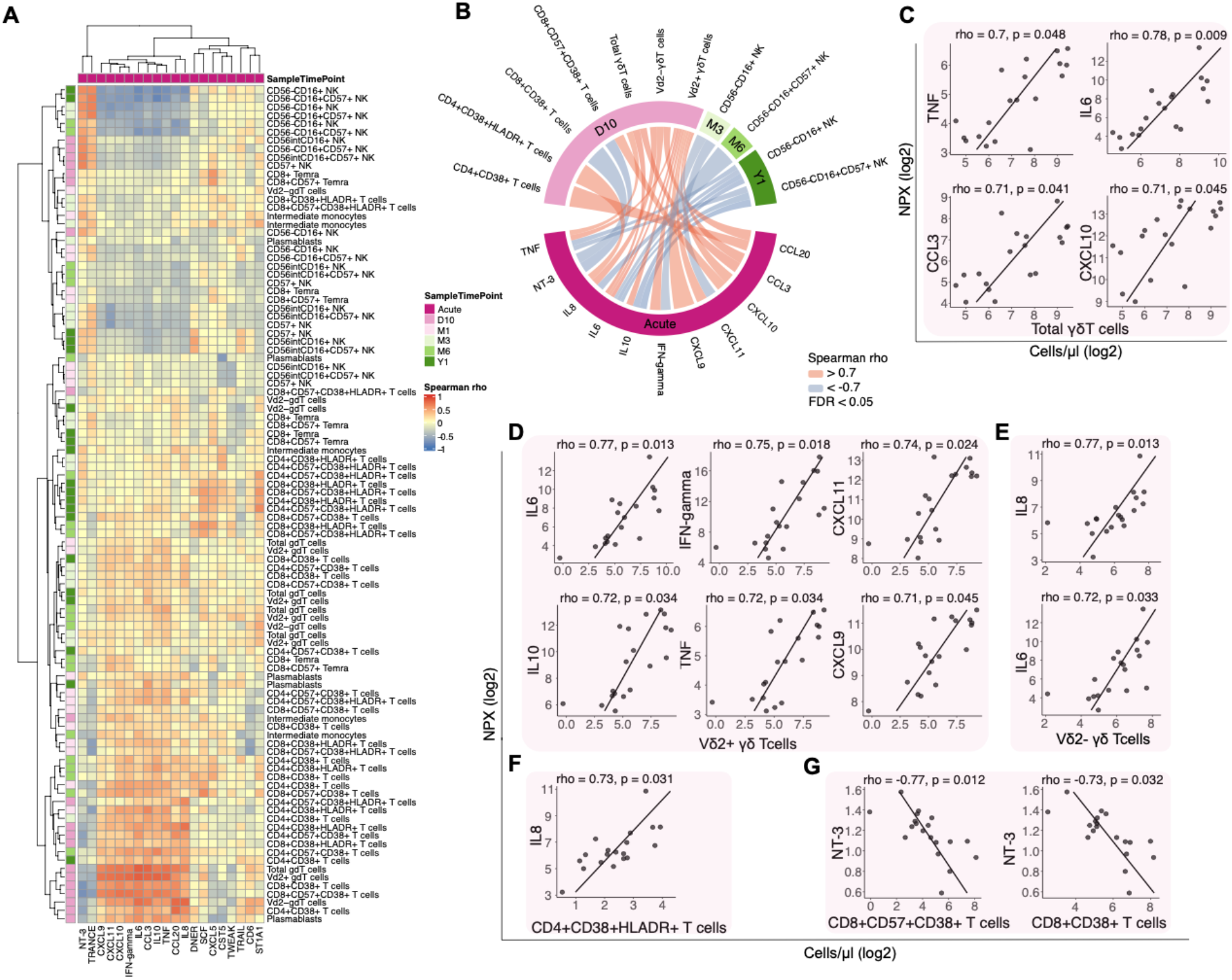
Associations of the acute phase plasma protein response with immune cell subsets. (**A**) Heatmap based on overall Spearman correlation (rho) acute phase plasma proteins and post-acute phase cell immune cell subsets. Blue and red colors symbolize positive and negative correlations, respectively. **(B)** Chord diagram based on strong (rho > |0.7|) and significant correlations values (after adjusted FDR < 0.05). (**C-G**) Scatter plots to show acute phase plasma protein levels and immune cell subset counts at day 10 post disease for significant correlations of **(C)** Total γδ T cells, **(D)** Vδ2^−^ γδ T cells, **(E)** Vδ2^+^ γδ T cells, **(F)** activated CD4+ T cells and **(G)** activated CD8+ T cells. All stated p-values are FDR corrected for multiple testing.

In particular, the numbers of γδ T cell subsets at the 10-day sample time-point were strongly associated with several pro-inflammatory cytokines at the acute time-point. The total level of γδ T cells positively correlated with TNF (rho = 0.704, p = 0.049), IL6 (rho = 0.781, p = 0.009), CCL3 (rho = 0.712, p = 0.041), and CXCL10 (rho = 0.707, p = 0.045) (Fig. 3C). Especially the Vδ2^+^ subset of γδ T cells has been shown to be important in the immune response to malaria parasites *(21–23)*. When investigating these subsets, we observed that the number of Vδ2^+^γδ T cells was positively correlated with levels of IL6 (rho = 0.767, p = 0.013), IFN-gamma (rho = 0.753, p = 0.018), CXCL11 (rho = 0.74, p = 0.024), IL10 (rho = 0.723, p = 0.034), TNF (rho = 0.724, p = 0.034), and CXCL9 (rho = 0.707, p = 0.045) (Fig. 3D). The number of Vδ2^−^ γδ T cells in contrast was only positively correlated with levels of IL8 (rho = 0.77, p = 0.013), and IL6 (rho = 0.72, p = 0.033) (Fig. 3E).

Acute phase levels of IL8 were also positively correlated with the number of activated (CD38^+^HLA-DR^+^) CD4^+^ T cells (rho = 0.728, p = 0.031) at the 10-day sample time-point. In contrast, neurotrophin-3 (NT-3) levels were negatively correlated with the number of activated CD8^+^ T cells in several different subsets (CD57^+^CD38^+^; rho = –0.77, p = 0.012 and CD57^+^CD38^+^HLA-DR^+^; rho = –0.726, p = 0.042) (Fig. 3F, G).

These results point towards a strong association of the levels of pro-inflammatory cytokines during the early acute phase and the size of immune cell subsets after the acute phase. The highest degree of association was identified for γδ T cells, and especially the Vδ2^+^ subset, which was strongly positively correlated with the levels of pro-inflammatory cytokines.

### Subhead 4: Impact of previous *P. falciparum* exposure on γδ T cell responses

To further characterize the association between acute phase cytokine levels and the γδ T cell response, we sought to examine if previous exposure to malaria parasites impacts this association. Overall, both groups show similar trajectories of their immune dynamics when plotting Factor1 values against time (Fig. 4A). However, primary infected individuals were associated with both higher and lower factor values during the acute phase and convalescent phase, respectively (Fig. S2C). This indicates differences in protein and immune cell subset magnitudes attributed to previous malaria.

**Fig. 4.**
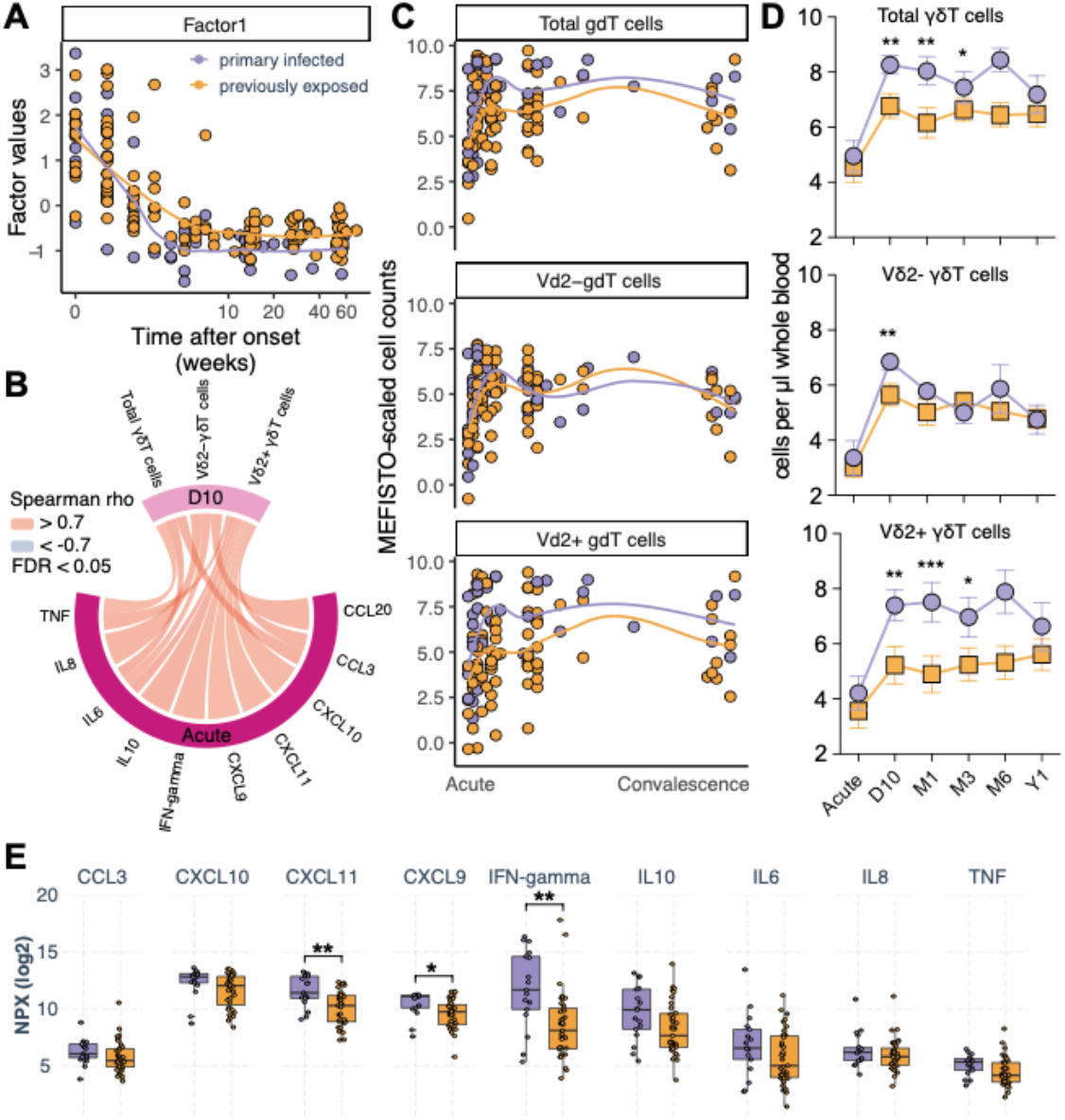
Impact of previous P. falciparum exposure on immune dynamics after clinical malaria. **(A)** Smooth fit of MEFISTO Factor1 values against time after symptom onset in weeks visualizing transition from acute to convalescence for primary infected (purple) and previously exposed (orange) individuals. **(B)** Associated inflammatory signature with γδ T cells based on spearman rank correlation. **(C)** γδ T cell dynamics over time after symptom onset. (**D**) Longitudinal γδ T cell levels. Statistical differences between groups for γδ T cell subtypes were assessed using linear mixed model fit with restricted maximum likelihood and t-tests with FDR correction for multiple testing. Error bars denote standard error of the mean. **(E)** Comparison between primary infected (purple) and previously exposed (orange) individuals for acute phase proteins significantly associated with γδ T cells at the acute time-point. Statistical differences between groups were assessed using the non-parametric Wilcoxon-test, FDR adjusted p-values to correct for multiple testing. *p < 0.05, **p < 0.01, ***p < 0.001.

Due to the strong correlation of acute phase proteins and γδ T cell subset numbers at day 10 (Fig. 4B, excerpt of Fig. 3B), we aimed to investigate the association further in the context of previous exposure.

When plotting γδ T cells against the time after symptom onset, we observed that Vδ2^+^γδ T cells expanded to a greater extent in primary infected individuals while Vδ2^−^ γδ T cells expanded similarly for both groups (Fig. 4C). We confirmed that especially the Vδ2^+^ subset was significantly more expanded after the acute time-point in primary infected individuals (Fig. 4D). All acute phase proteins that correlated with Vδ2^+^ γδ T cells (Fig. 4B) followed a similar trend, although only IFN-gamma, CXCL9 and CXCL11 were significantly higher in primary infected compared to previously exposed individuals (Fig. 4E).

Here, we could show that primary infected individuals have significantly higher levels of pro-inflammatory cytokines at the acute phase, as well as a significantly larger expansion of Vδ2^+^ γδ T cells after the acute phase compared with individuals previously exposed to malaria.

### Subhead 5: Effect of previous malaria exposure on Vδ2^+^ γδ T cell characteristics

The Vδ2^+^ subset of γδ T cells have been shown to play an important role in combating blood stage malaria, via TCR-induced cytotoxicity and CD16 mediated phagocytosis *(24, 25)*. We therefore focused on characterizing the CD16^+^Vδ2^+^γδ T cell response further using linear-mixed effects models to assess the impact of previous exposure to parasites on γδ T cell functional markers.

First, we assessed CD16 expression among γδ T cells. The expression was similar between primary infected and previously exposed individuals for total and Vδ2 positive and negative subsets (Fig. 5A). The frequency of CD16^+^ cells increased somewhat in both groups between the acute timepoint and 10 days, for Vδ2^−^ cells. A similar expansion was also observed for the Vδ2^+^ cells, but only in primary infected individuals (Fig. 5A), potentially reflecting an upregulation in response to infection. To further assess if the Vδ2^+^γδ T cells responded to the infection, we next investigated the activation status of the CD16^+^Vδ2^+^γδ T cells. Approximately 60% of all Vδ2^+^γδ T cells had upregulated the activation marker CD38 at the acute infection. The primary infected individuals then retained a significantly higher frequency of CD38^+^ cells at the day 10 time-point after which the levels reduced over 3-6 months until reaching baseline levels (Fig. 5B). The increased activation-status in primary infected individuals was also reflected by a significantly higher co-expression of CD38 and HLA-DR at the acute time-point (Fig. 5B).

**Fig. 5.**
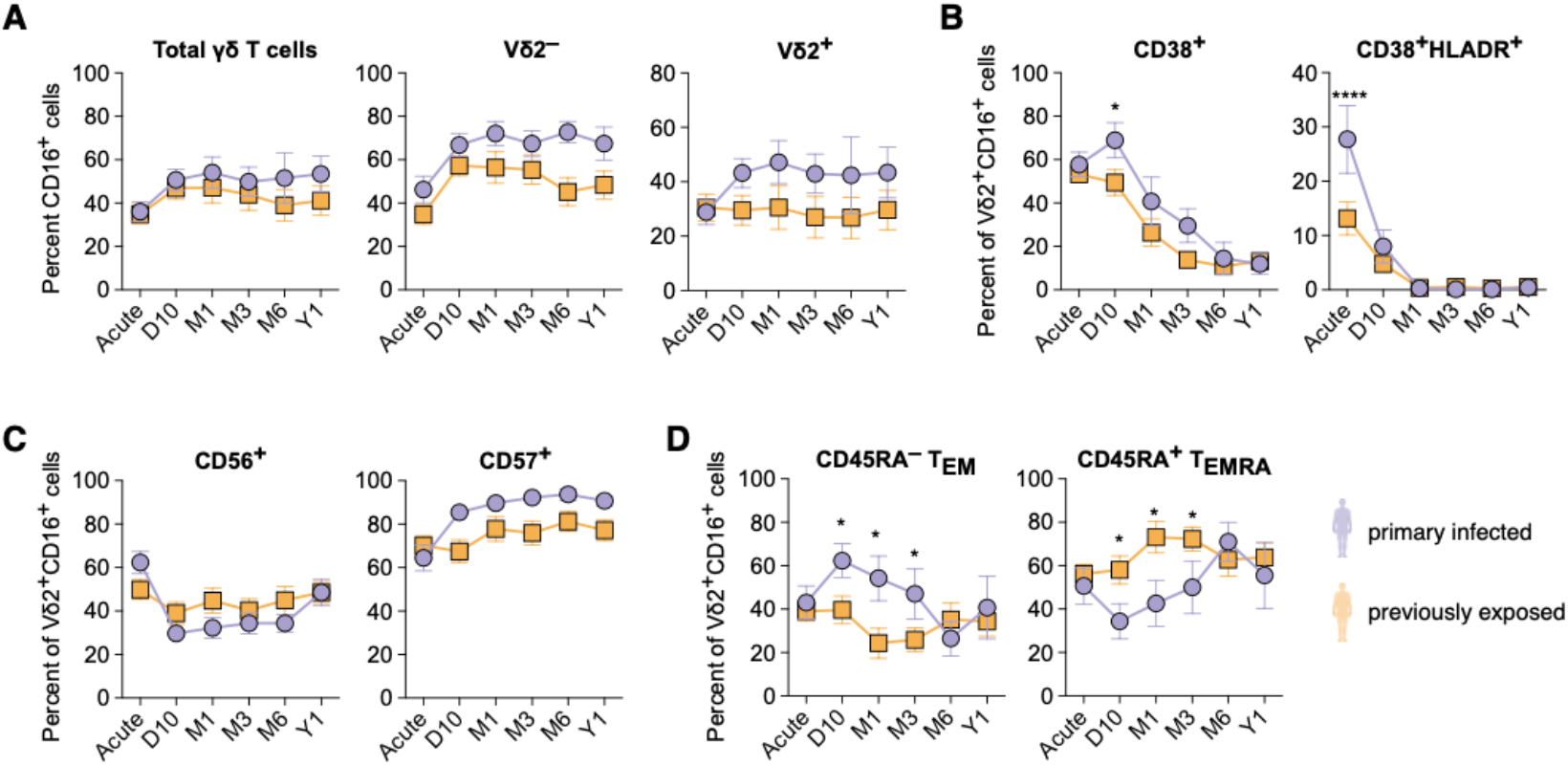
Previous P. falciparum exposure impacts γδ T cell expansion. Linear mixed effect modelling by maximum likelihood for γδ T cell frequencies for primary infected (purple) and previously exposed (orange) individuals, error bars denote the standard error of the mean. (**A)** Percentage of CD16^+^ cells of total γδ T cells, Vδ2^−^ or Vδ2^+^ γδ T cells. **(B)** Percent of Vδ2^+^ CD16^+^γδ T cells expressing activation markers CD38 alone or together with HLA-DR. (**C**) Percent of Vδ2^+^CD16^+^γδ T cells expressing CD56^+^ or CD57^+^ **(D)** Vδ2^+^γδ T cells effector memory (CD45RA^−^) or T_EMRA_ (CD45RA^+^). Difference in cell frequency at each time point was evaluated by repeated t-tests with FDR correction. *p < 0.05, ****p < 0.0001.

We also assessed the expression of CD56 and CD57, associated with NK cell cytotoxicity *(26)* and replicative senescence *(27)*, respectively. Although there were no differences between the groups, there were some changes in cells expressing the markers over time, such as a temporary reduction in the frequency of CD56^+^ cells after the acute infection, while CD57^+^ cells increased over time (Fig. 5C). These effects were especially observed in primary infected individuals, potentially reflecting the stronger γδ T cell response in these individuals.

To further determine which subsets of Vδ2^+^γδ T cells that responded during infection, we assessed the levels of CCR7 and CD45RA to identify different naïve or effector populations *(28)*. Almost all CD16^+^ Vδ2^+^ γδ T cells were negative for CCR7 (median 98 % over all time-points), indicating that these cells display an effector-phenotype. During the acute phase of the response, both groups displayed similar distribution of effector memory (T_EM_, CD45RA^−^) and T_EMRA_ (CD45RA^+^) γδ T cells (Fig. 5D). However, as the number of Vδ2^+^γδ T cells expanded in primary infected individuals the frequency of effector memory cells significantly increased, while the frequency in previously exposed individuals was relatively stable over time. This suggests that it was primarily effector memory Vδ2^+^γδ T cells that expanded after the acute infection.

Collectively these results show that despite Vδ2^+^ γδ T cells only expanding in primary infected individuals, the cells became activated in both groups after infection. The level of activation and subsequent changes in host effector and differentiation markers were however different with a more robust effect in primary infected individuals.

### Subhead 6: γδ T cell function is retained while expansion is affected due to previous malaria exposure

It has been described that especially Vδ2^+^γδ T cells respond strongly during malaria but that this effect is reduced upon subsequent infections *(29)*. This pattern is supported by our data, as the Vδ2^+^γδ T cell activation and especially expansion were reduced in previously exposed compared to primary infected individuals (Fig. 4D). Previous descriptions indicate that the reduced responsiveness can be due to changes in the imprinting through changes in the methylation patterns *(30, 31)*, although it remains unclear if such imprinting can last for this long. To assess if the function of the γδ T cells in our cohorts were inherently affected in their capacity to respond to stimulation, we performed blinded cultures of PBMCs from primary infected and previously exposed individuals. We stimulated the cells with PMA and ionomycin that together bypass the T cell receptor complex and measured production of TNFa, IL17, and IL10. PMA and ionomycin stimulation led to increased production of TNFa, but not IL17 or IL10 (Fig. 6A). Comparing the response of Vδ2^+^ and Vδ2^−^γδ T cells, the Vδ2^+^ cells responded with more TNFa production (Fig. 6B), consistent with a proinflammatory effector function. For Vδ2^−^ γδ T cells, the production of TNFa was similar between the acute and convalescent time-point and healthy controls. For Vδ2^+^γδ T cells, however, the frequency of TNFa producing cells was reduced during the acute response, but then recovered at the convalescent phase, for which frequencies were similar as in healthy controls (Fig. 6C). Further separating the Vδ2^+^ and Vδ2^−^ subsets into primary infected and previously exposed individuals showed no differences between the groups for Vδ2^−^ cells, while primary infected individuals responded with lower numbers of TNFa-producing Vδ2^+^ cells at the acute time-point (Fig. 6D), potentially reflecting a larger proportion of already activated cells being restimulated.

**Fig. 6.**
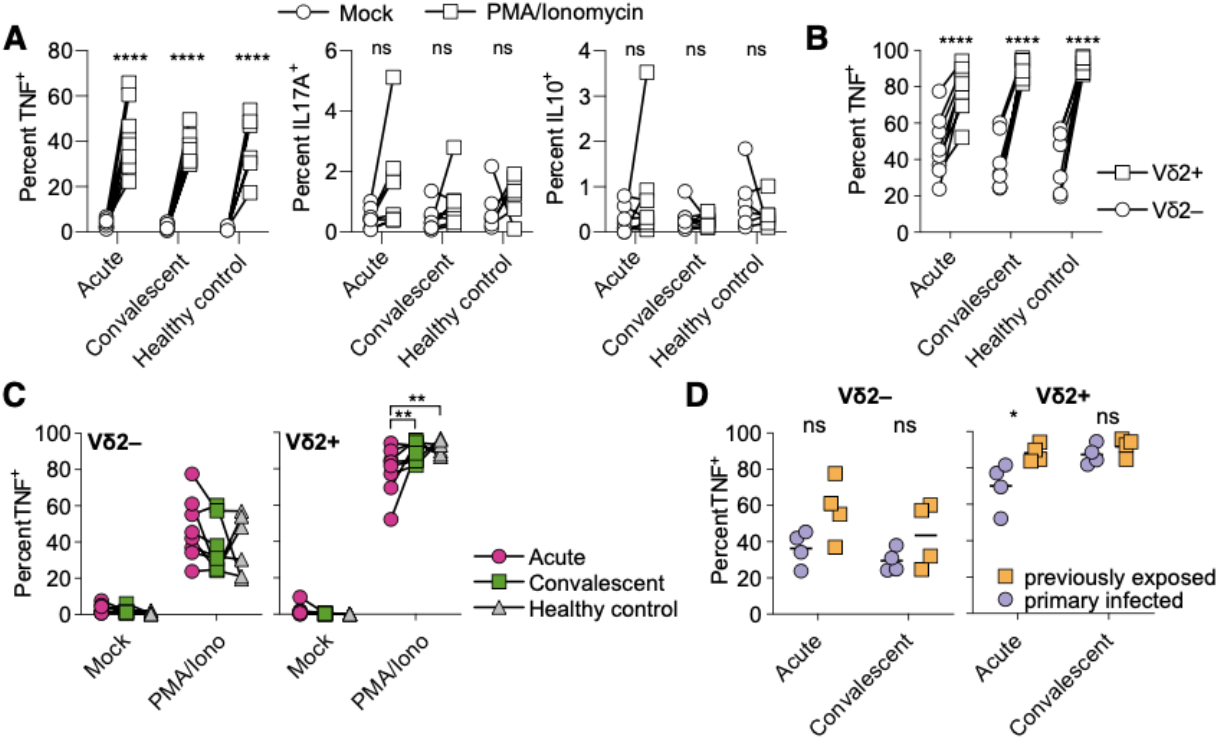
γδ T cell function in response to restimulation. (**A**) PBMCs from donors with primary infection (n = 4) and previous parasite exposure (n = 4) at the acute and convalescent time-point (6-12 months after infection) and healthy controls (n = 6) were stimulated with PMA and ionomycin (open boxes) or left unstimulated (Mock, open circles) for 5 hours. Frequencies of TNFa, IL17A, and IL10-producing cells were then measured. (**B**) TNF^+^ cells were compared between Vδ2^+^ (open boxes) and Vδ2^−^ (open circles) cell subsets. (**C**) Comparison of TNF^+^frequencies in Vδ2^−^ and Vδ2^+^ cell subsets at the acute time-point (pink circles), convalescent time-point (green boxes) and healthy controls (grey triangles). (**D**) Comparison of TNF^+^ Vδ2^−^ and Vδ2^+^ cells between primary infected (purple circles) and previously exposed (orange boxes) individuals. Statistical comparisons for A-C were done using a matched pair two-way ANOVA followed by Tukey’s post-hoc test, while statistics in D was evaluated by two-way ANOVA followed by Sidak’s post-hoc test. *p < 0.05, **p < 0.01, ****p < 0.0001, ns = not significant.

In summary, Vδ2^+^γδ T cells from previously exposed individuals do not display apparent intrinsic dysfunctionality upon *ex vivo* reactivation. This further supports a role for extrinsic factors regulating Vδ2^+^γδ T cell activation and expansion in malaria.

### Subhead 7: Acute malaria specific IgG3 levels are associated with the level of inflammation and γδ T cell expansion

A long-lived memory imprint to pathogenic encounter is in general mediated by the adaptive immune system. Here, we report that previously exposed individuals without re-exposure to parasites for many years (average 11.5 years since leaving an endemic region), at the time of a new acute infection respond with a reduced pro-inflammatory cytokine response upon re-infection as well as reduced Vδ2^+^γδ T cell expansion. Hence, we hypothesized that the reduced pro-inflammatory response and γδ T cell expansion could be mediated by existing long-lived adaptive humoral immune memory responses.

To investigate this, we repurposed a previously published dataset, investigating the immunoglobulin G subclass response (IgG1-4) to five malaria *P. falciparum* blood-stage antigens (AMA1, MSP1, MSP2, MSP3, and RH5) in 51 matching individuals from the same cohort *(15)*. We established an IgG subclass and time point specific score, the cumulative response score (CRS) that summarizes the responses to the separately measured malaria specific antigens from the previous study (Fig. S5). The CRS dynamics correspond to the average breadth of the antibody response for total IgG and IgG subclasses (Fig. 7A).

**Fig. 7.**
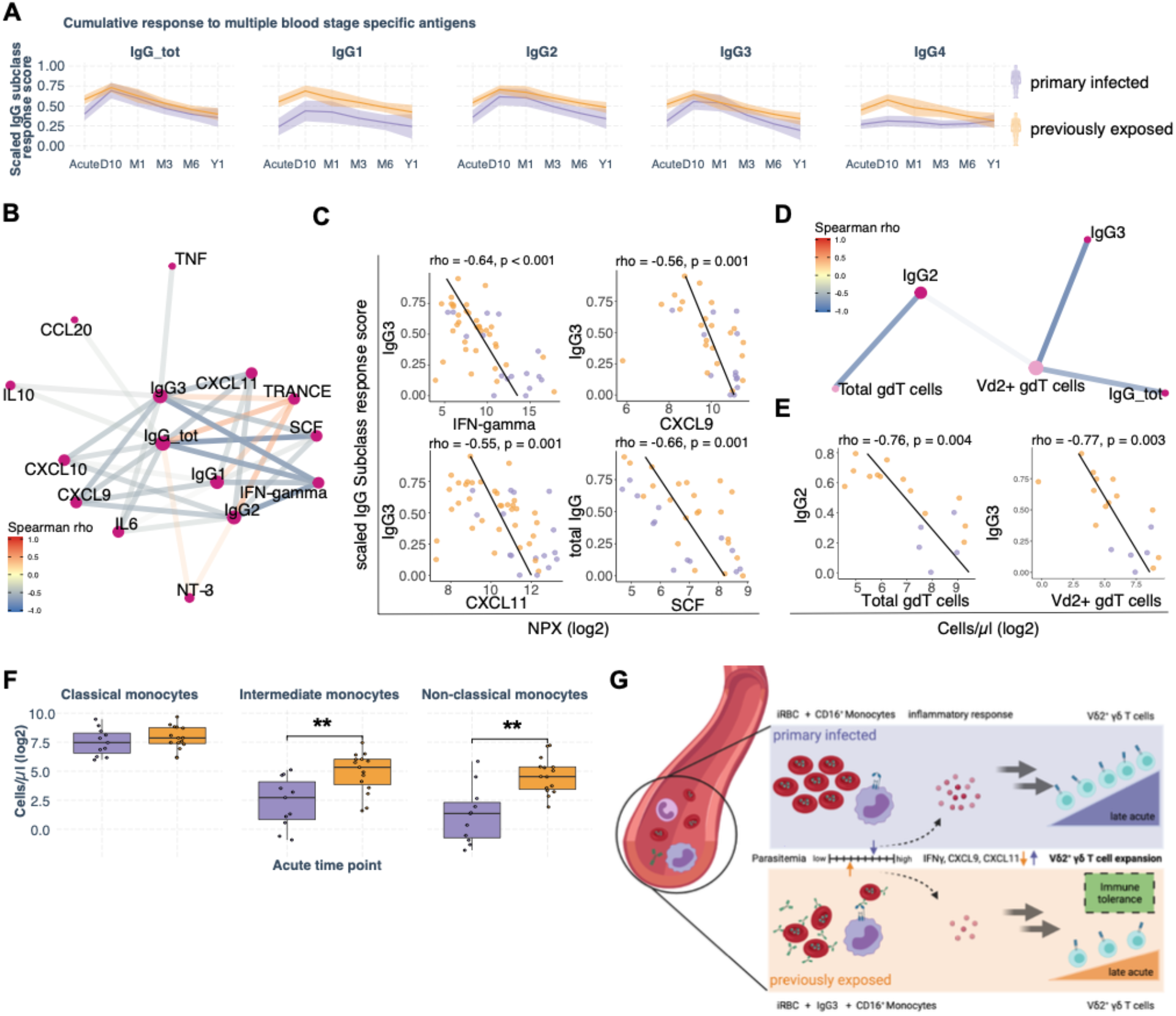
Adaptive response due to prior malaria exposure linked to dampened immune inflammation. (**A**) Cumulative IgG subclass response score (CRS) to 5 blood-stage antigens from Yman et al., BMC Medicine, 2019, over sample time points for primary infected (purple) and previously exposed (orange) individuals (n = 52), shaded area denotes the 95% confidence interval. **(B)** Correlation network of plasma proteins and IgG subclass CRS values at the acute time-point, FDR corrected p < 0.05 **(C)** Scatter plots for significant correlations, FDR adjusted p-values are stated. **(D)** Correlation network of acute IgG subclass CRS values and immune cell subsets at the 10-day sample time point, FDR corrected p < 0.05, rho < –0.7. **(E)** Scatter plots for significant correlations, FDR corrected p < 0.05. **(F)** Cell counts of monocyte subsets at the acute time point. Statistical differences between groups were assessed using the non-parametric Wilcoxon-test, FDR adjusted p-values to correct for multiple testing. *p < 0.05, **p < 0.01 **(G)** Schematic model to explain improved disease tolerance in previously exposed individuals.

Spearman rank correlation showed that levels of several acute pro-inflammatory cytokines were strongly inversely correlated with IgG subclass CRS (Fig.7B), in particularly IgG3 and IFN-gamma levels (Fig. 7C). The IgG2 and IgG3 subclass CRSs at the acute infection were also negatively correlated with day 10 total and Vδ2^+^γδ T cell numbers, respectively (Fig. 7D), with an especially strong negative correlation between IgG3 levels and Vδ2^+^γδ T cell numbers (Fig. 7E).

Cytophilic immunoglobulins such as IgG3 can interact with CD16 expressing immune cells, including γδ T cells, and intermediate (CD16^+^CD14^high^) and non-classical (CD16^+^CD14^l0^) monocytes. Interestingly, we observed higher numbers of intermediate and non-classical monocytes at the acute time point for previously exposed compared to primary infected individuals (Fig. 7F). These observations, together with an increased IgG3 response, points towards an antibody-phagocytosis mediated mechanism that could partly explain the lower parasitemia observed in individuals with prior exposure to malaria parasites (Fig. 1E). To assess if this could be the case, we used linear regression to determine which associated combinations of immune factors that best explain the difference in expansion of Vδ2^+^γδ T cells for the two exposure groups. Using Directed Acyclic Graphs (DAG) *(32)*, we determined the minimal sufficient adjustment sets of covariates for estimating the total effect of a given associated component on acute IFN-gamma levels and day 10 Vδ2^+^γδ T cell numbers, respectively (described in Supplementary Methods and Fig. S7). Comparing the reference model with exposure group as explanatory variable to the DAG-based adjusted immune response component model, we concluded that the IFN-gamma response was not directly explained by the parasitemia (Fig. S7A). In contrast, differences in Vδ2^+^γδ T cell expansion was partly explained by parasitemia, but more directly by cytophilic antibody levels and CD16^+^ intermediate monocytes (Fig. S7B-C). This suggests a potentially more direct antibody-mediated effect by regulating cytokine levels. Based on these results we suggest a working model where early production of parasite-specific antibodies interact with innate immune cells to regulate parasitemia and cytokine levels, which in turn control subsequent γδ T cell expansion (Figure 7G).

## DISCUSSION

Immunity to malaria is slow to develop. It has been proposed that this is partly due to immune perturbations during infection *(41)*. Understanding how the immune system is affected during natural infection with malaria parasites is important to identify the potential role of individual immune mediators in this process. Here, we used systems immunology to comprehensively investigate the immune landscape after natural malaria and over one year after infection in the absence of re-exposure to parasites. By sampling each individual over time, we could reduce potential noise associated with inter-individual variability in their immune response to the infection. Overall, the integrated immune landscape identified the acute phase followed by a transitional phase leading into convalescence up to one year after disease. The acute phase was characterized by activated CD4^+^ and CD8^+^ T cells and high levels of inflammatory cytokines and chemokines, consistent with previous literature *(2, 33)*. Here, a comprehensive analysis of immune components over time revealed that acute inflammatory response signature associated with long-term changes to cellular immune response.

The composition of our cohort, consisting of both primary infected and previously exposed individuals, allowed us to study the impact of memory on the acute response and its associations with the transition towards immunological convalescence. We could show that previously exposed individuals produce lower levels of pro-inflammatory cytokines during natural *P. falciparum* infection. Similar findings have been observed at the blood transcriptome level by Tran et al., where Malian adults exhibited a dampened inflammatory response compared to first time CHMI-infected Dutch adults *(34)*. Further, a study from a cohort of children in Uganda showed that older children have lower levels of serum cytokines during acute malaria compared to younger children *(35)*. These previous studies, together with our findings of reduced pro-inflammatory response confirms that previous malaria exposure, possibly cumulative, induces a form of tolerance response. Of note, all individuals in our cohort experienced clinical malaria, suggesting that the level of tolerance could be dependent on the amount of prior exposure to the parasite. However, the individuals with previous exposure had left malaria endemic areas on average 11.5 years earlier before being reinfected/experiencing a new episode of acute *P. falciparum* malaria after visiting endemic area, indicating that the mechanism is significantly long-lived.

Antibodies are highly associated with protection from malaria and a key building block of naturally acquired immunity. We have recently reported that previously exposed individuals from our cohort responded to infection by producing high levels of *P. falciparum*-specific IgG, particularly of the cytophilic subclasses IgG1 and IgG3 *(15)*. Memory B cells and long-lived plasma cells constitute a likely source of the increased parasite specific antibodies *(15, 36)*, as memory can be maintained for up to 16 years without re-exposure in a different cohort of travelers *(37)*. Re-purposing the previously published antibody data *(15)* enabled us to link the effect of adaptive responses, in the form of an increased cytophilic antibodies, to enhanced functionality of the innate response. As reported here, previously exposed individuals responded to malaria with an overall dampened inflammatory response but significantly higher numbers of intermediate monocytes *(38)*. Their expression of CD16 (FcγRIII), a receptor for cytophilic antibodies of the subclasses IgG1 and IgG3, can mediate parasite clearance through antibody-dependent phagocytosis (ADP) and/or antibody-dependent cellular cytotoxicity (ADCC) *(39)*. Indeed, higher numbers of CD16^+^monocytes and a stronger IgG3 response against blood stage antigens indicate an interplay which could promote lower levels of parasitemia, compared to individuals without previous exposure. Supporting such a mechanisms, vaccination with RTS,S/AS01 leads to CD16-mediated phagocytosis associated with protection *(40)*.

We propose that antibody-mediated parasite control is an important component in shaping the pro-inflammatory cytokine milieu towards a reduced pro-inflammatory response which could be important for the development of immune tolerance *(41)*. Complementary to the described adaptive-innate interplay, several recent studies suggest that monocytes are modulated towards a tolerance phenotype *(5, 41)*. The mechanism of this imprint remains unclear, but a rodent model point towards transcriptomic reprogramming of monocytes in the spleen *(5)*, while data from *in-vitro* stimulated monocytes from a cohort in Mali point towards epigenetic reprogramming of myeloid progenitor cells in the bone marrow *(41)*. However, to our knowledge, no study has shown that epigenetic remodeling can remain and affect innate cells at re-stimulation for that long. An alternative hypothesis, that overcomes the fact that monocytes have a short lifespan *(42)*, is that they are remodeled in their function due to constant stimuli by remaining hemozoin in the spleen *(5, 43, 44)*.

Both hypotheses of epigenetic imprinting and through splenic remodeling need further investigation. In our cohort, where study participants moved away from malaria endemic areas 10 years prior, we propose an additional mechanism responsible for the tolerogenic response via the interplay of innate and adaptive immunity. We suggest that tolerance develops over several exposures to parasites and further that it is sustained long-term through adaptive memory responses. However, these above hypotheses are not mutually exclusive and could potentially synergize.

In agreement with previous reports, we observed expanding γδ T cells after the initial inflammatory response in response to the infection *(45, 46)*. γδ T cells are suggested to have an important role in the control of malaria *(29)* and direct anti-parasite functions during the blood stage have been reported *(24)*. In this study, we observed that the acute phase inflammatory response was positively associated with expansion of Vδ2^+^ γδ T cells, and further that this was strongly impacted by previous malaria exposure, resulting in a dampened inflammatory response and less γδ T cell expansion. Interestingly, this effect was primarily observed for the subset of Vδ2^+^ and not Vδ2^−^ γδ T cells. The strong association could indicate that the pro-inflammatory response directly or indirectly shapes the subsequent γδ T cell expansion and capacity to adapt to prolonged and high levels of blood stage parasitemia.

Vδ2^+^ γδ T cells are known to activate and expand during a primary *P. falciparum* infection in response to malaria phosphoantigens and that their activity is modulated upon subsequent infections *(47)*. Given the recently described role of Vδ2^+^γδ T cells in antiparasitic activities via antibody-CD16 dependent phagocytosis of infected erythrocytes and cytotoxicity *(24, 25)*, associations to possible activity modulating factors are of interest. The benefit of reduced expansion of γδ T cells due to a dampened inflammatory response could be to reduce overall inflammatory responses, which are otherwise potentially detrimental to the host *(48)*. This is a common hypothesis as the γδ T cells are known to acquire a dysfunctional or tolerance phenotype over time with repeated episodes of malaria *(23)*. This reduction in γδ T cell effector function was associated with continuous malaria exposure *(23)*, although it remains unclear what the underlaying tolerance mechanism is, and how long-lived such an effect could be. To assess if the reduced activation and expansion of Vδ2^+^γδ T cells could be due to an inherent inability of the γδ T cells to respond, we re-stimulated cells from primary infected and previously exposed individuals. However, we did not observe any reduced response that could have been associated with intrinsic down-regulation of effector functions. Based on this, we instead hypothesized that the regulation of Vδ2^+^γδ T cell activation and expansion was likely due to extrinsic mechanisms.

Among the cytokines and chemokines associated with Vδ2^+^γδ T cell expansion, we found IFN-gamma and CXCL11 significantly reduced in previously exposed individuals. CXCL11 is chemotactic for activated T cells, and it was reported that individuals with asymptomatic *P. falciparum* malaria had lower levels in an endemic setting *(49)*. IFN-gamma and the IFN-gamma inducible chemokine CXCL9 are associated in a different context with regulatory crosstalk of pro-inflammatory γδ T cell effects *(50)*, which could explain the association of these cytokines with subsequent expansion. Consistent with these observations, both IFN-gamma and CXCL11 were reduced due to previous parasite exposure in a mouse model, suggesting some type of memory response *(5)*.

In summary, our results, together with previous research in the field, supports a model where early control of host self-damage through increased tolerance is needed until a more broad, diverse, and protective antibody repertoire is achieved (see Figure 7G). The tolerance response (dampened pro-inflammatory response and reduced Vδ2^+^γδ T cell expansion) could be mediated in two ways: 1) an expanding antibody repertoire that enhances CD16^+^ mediated phagocytosis and effector functions, direct parasite neutralization and blocking further RBC invasion, rosetting and sequestration by merozoites, and 2) training/priming during previous malaria episodes via epigenetic or transcriptional remodeling of the monocyte population, potentially leading to rapid generation of CD16-expressing monocytes with enhanced antibody effector functions.

From an evolutionary perspective, this type of dampened inflammatory response could be important to generate an efficient B cell response, as recently shown in an influenza vaccine study *(51)* and further supported by mouse models of malaria *(52)*. Although further studies are needed to evaluate these mechanisms on a cellular and molecular level, the comprehensive systems levels analysis performed here provides a dynamic description for how proteins, cells, and antibodies, interact in the host response to malaria during and after primary and repeated infection.

## MATERIALS AND METHODS

### Study design/cohort

A prospective study enrolling adult patients diagnosed with *P. falciparum* malaria at Karolinska University Hospital in Stockholm, Sweden. Fifty-three patients that were admitted with acute *P. falciparum* infection between 2011 and 2017, and where both cells and plasma was frozen were included in the current study. We stratified the patients based on previous exposure to compare the immune responses in malaria-naive individuals of primarily Swedish origin, who contracted malaria for the first time (n = 17, denoted as primary infected), with individuals originating from malaria-endemic areas in Sub-Saharan Africa and reporting previous malaria (n = 36, denoted as previously exposed). Patients were invited for sampling at the time of malaria diagnosis (Acute) and then at approximately 10 days (D10), 1 month (M1), 3 months (M3), 6 months (M6), and 12 months (Y1) (see Fig.1A). On each sampling, venous blood was collected for plasma and serum isolation, PBMC preparation and blood chemistry. Peripheral blood mononuclear cells (PBMCs) were isolated using Ficoll-Paque density gradient separation, resuspended in 90 % fetal calf serum supplemented with 10% DMSO, and stored at - 150°C. Clinical data was extracted from medical records, and a questionnaire relating to the participant’s health status, previous traveling, and malaria exposure was filled in by all participants (Table S1).

Part of the cohort has been described previously for clinical *(53)*, parasitological *(13)* and immunological aspects *(15, 16, 54)*.

In addition, we profiled peripheral blood samples of eight healthy controls to compare relative measures of protein expression and cell populations with healthy/normal values (Figure S2).

### Clinical diagnostics

*P. falciparum* parasites were detected and enumerated by light microscopy of Fields stained thick and thin blood smears at the Department of Clinical Microbiology at Karolinska University Hospital in addition to PCR as described previously *(13)*. Leukocyte, neutrophil, monocyte, and platelet cell differential counts were performed at the Department of Clinical Chemistry at Karolinska University Hospital.

### Data acquisition

#### Immune cell phenotyping

Immune cells were phenotyped by Flow cytometry (LSRFortessa, BD) and a panel of 17 antibodies covering major immune cell populations and subpopulations (Fig. S1) for 137 samples. Frozen PBMCs were thawed in a 37°C water bath and mixed with 1 equal volume cold Iscove’s Modified Dulbecco’s Medium supplemented with L-glutamine (2 mM), penicillin (100 U/ml), streptomycin (100 μg/ml), and 10 % heat-inactivated fetal bovine serum (all from Thermo Fischer Scientific). Cells were then rested 20 minutes on ice before being washed twice in DPBS lacking magnesium or calcium. After washing, the cells were incubated with Aqua Live/Dead stain (Thermo Fisher Scientific) for 20 minutes followed by further washing in DPBS supplemented with 2% FBS. The cells were then incubated for 20 minutes on ice, in 2 steps with 2 washes in between, using an antibody mix targeting surface antigens (Table S2). After staining, the cells were washed twice in DPBS with 2% FBS before acquisition on a 5-laser BD LSRFortessa flow cytometer. Gating was done with FlowJo X software version 10.4.2, with the gating strategy shown in Figure S1.

#### Plasma protein profiling

Plasma proteins relevant for the immune response were profiled using the Proximity Extension Assay (PEA) (Olink Proteomics, Sweden) to analyze 171 samples of 53 individuals using Olink Target 96 Inflammation panel (92-plex). The method has been described previously *(17)*. Briefly, paired oligonucleotide-coupled antibodies bind to target proteins, leading to hybridization of the oligonucleotides when the antibody-pair is in close proximity, forming a PCR template for real-time PCR detection. Resulting data is normalized using assay internal controls and transformed into Normalized Protein eXpression (NPX) values, representing an arbitrary relative quantification unit on log2 scale. After quality control and removal of markers with a missing frequency greater than 50%, 74 proteins remained for downstream analysis.

#### *In vitro* stimulation for detection of γδ T cells cytokine production

For intracellular staining, PBMCs were thawed, washed, resuspended in complete RMPI media supplemented with 10% FCS and then incubated at 37°C over-night. The cells were then stimulated with PMA (0.4 μM) and ionomycin (13 μM) per 10^6^ cells, (Nordic Biosite AB) for 5 hours. Golgi plug (BD Biosciences) was added after the first hour of stimulation. Activated cells were subsequently stained with antibodies targeting the surface markers CD3, γδ TCR, and Vδ2, according to manufacturers’ instruction. Cells were then fixed and permeabilized using the BD Cytofix/Cytoperm reagents (BD Biosciences) and stained with intracellular antibodies for TNF, IL17A, and IL10 (Table S2). For the exclusion of dead cells, LIVE/DEAD aqua fixable viability staining kit (Invitrogen) was used. Cells were acquired using a BD LSR Fortessa BD (BD Biosciences) and the data analyzed using FlowJo Version 7.6 (FlowJo, Ashland, OH).

### Bioinformatics

#### Immune cell data normalization

Prior to bioinformatic analysis, cell proportions from manually gates (Fig. S1) were normalized to the number of cells per 1000 live cells. Subsequently, we adjusted the experimentally obtained live cell counts with lymphocyte and monocyte counts from clinical blood chemistry counts for each individual and timepoint to obtain cells per microliter blood values (Fig. S5A). Monocyte gates containing values (Fig. S1A) were normalized according to:

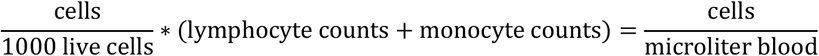

Values of pure lymphocyte containing gates (Fig. S1B) were normalized according to:

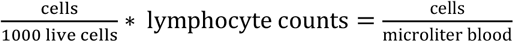

To standardize the cells-per-microliter values and to normalize the data set for extreme values, we performed log2 transformation. Standardized data was comparable to not-standardized data (Fig. S5B).

#### Antibody cumulative response score (CRS) calculation

A previous study on individuals of the traveler cohort investigated the immunoglobulin G subclass response (IgG1-4) s to five malaria *P. falciparum-blood-stage* antigens (AMA1, MSP1, MSP2, MSP3, RH5) in 52 this current study matching cohort individuals *(15)*. For the current study, we though to look at the overall IgG subclass response to merozoite antigens instead of specific antigens. To summarize the adjusted signals, measured for each antigen and each subclass, we rank normalized the signals to achieve normal distribution for each antigen (Fig. S5A). Normally distributed antigen response values were then summarized for each IgG subclass and time point, creating the cumulative response score (CRS), presenting an average breadth value of the antibody response for IgG subclasses (Fig. S5B).

### Statistical analysis and visualization

#### Multi-Omics-Factor Analysis

Multi-Omics Factor analysis (MOFA) is a powerful tool for omics integration. It reduces high-dimensional data into a few latent factors by capturing multi-dataset spanning variance in these factors. The unsupervised nature of MOFA+ allows the model to capture both biological and technical variability in the low-dimensional factors space *(18, 19)*. A recent development (MEFISTO) in addition allows for multi-omics integration of data with continues structures given by temporal relationships*(20)*. Here, we utilized MEFISTO to analyze the underlying key drivers for the time course of one year after infection on our cohort of returning travelers with *P. falciparum* malaria.

For the data-driven-integration approach, we used 15,396 immune feature data points out of 23,130 possible. We included manually gated immune cell subset (features = 44) data and highly multiplexed plasma protein data (features = 60) and including the time after symptom onset (0-70 weeks) for each sample to disentangle time dependent variation on a systems-level.

To standardize the actual time covariate of each sample, we used the time after reported symptom onset for each sample as a temporal covariate for the model. Moreover, we assigned the time-associated samples to groups of primary infected (n = 57) and previously exposed (n = 125) to account for previous *P. falciparum* exposure. Immune parameters without temporal variation were excluded prior to model training.

We trained the MEFISTO model using default model options, but adjusted training options (drop-factor-threshold = 0.05; maxiter = 10,000; convergence_mode = slow). To align the covariates across groups, warping was set “TRUE” and “primary infected” was set as warping reference group. The optimal model was determined by the MOFA+ function run_mofa() with the setting “use_basilisk = T”.

All data wrangling, analysis and visualization was done using R (www.r-project.org) using the tidyverse package *(55)*. Spearman correlation analysis with FDR/BH correction for multiple testing *(56)* was done using the correlation package *(57)*. If not stated otherwise, non-parametric data distribution was assumed, and statistical difference was assessed using unpaired Wilcox Test from rstatix (https://github.com/kassambara/rstatix).

Results were visualized using r packages circlize *(58)*, complexHeatmap *(59)* and ggpubr (https://github.com/kassambara/ggpubr9). Linear-mixed-effect models, with subjectID as random effect, were fitted in GraphPad prism version 9.1.2 using restricted maximum likelihood followed by t-tests using predicted least squares means with Benjamini and Hochberg FDR correction for multiple testing *(56)*.

## Supplementary Materials

Table S1. Descriptive statistics prospective cohort of returning travelers.

Table S2. Staining panel for flow cytometry

Fig. S1. Flow cytometry manual gating strategy - T cell and B cell panel.

Fig. S2. MEFISTO model and cohort internal feature evaluation.

Fig. S3. Cohort external evaluation - comparison of plasma protein and PBMC profiles at Acute and Y1 compared to healthy controls.

Fig. S4. Comparison of MEFISTO Factor1 values for exposure groups on time points.

Fig. S5. Cumulative Response Score for malaria antigen specific IgG subclass response of Yman et al 2019 data set.

Fig. S6. Leukocyte count adjusted cell counts.

Fig. S7. Directed Acyclic Graphs - DAGs and linear regression

## Acknowledgments

We would firstly like to acknowledge the contribution of all the study participants. We would also like to thank all involved clinicians, nurses especially Irene Nordling and Debbie Ribjer for sampling and Fariba Foroogh for sample processing and organization. Further, we would like to thank the team from the Translational Plasma Profile Facility at SciLifeLab for support and the generation of data for this project. Figure 1A and 7G were created with BioRender.com.

## Funding

The Swedish Research Council grant 2019-01940 (CS)

Magnus Bergvall foundation grant 2017-02043 and 2018-02656 (CS)

Åke Wiberg foundation grant M18-0076 (CS)

Swedish Research Council 2015-02977, 2018-02688 and 2018-04468 (AF)

Region Stockholm 20150135 and 20180409 (AF)

Marianne and Marcus Wallenberg Foundation (AF)

## Author contributions

Conceptualization: CS, MJL, AF

Methodology: CS, MJL, NK, SA

Investigation: CS, MJL, KS, VY, NK, SA, DFP, AF

Visualization: CS, MJL

Funding acquisition: CS, AF

Project administration: CS

Supervision: CS, AF

Writing – original draft: CS, MJL

Writing – review & editing: CS, MJL, AF, KS, VY, NK, SA, DFP

## Data and materials availability

The flow cytometry data and plasma protein data (Olink Inflammation panel) generated during this study are available upon request. Code used in the analyzes is available in repository: https://github.com/LautenbachMJ/MalariaTraveller.

