## Supplementary material for "Systems analysis shows a role of cytophilic antibodies in shaping innate tolerance to malaria"

**Supplementary Materials**

Materials and Methods

Table S1. Descriptive statistics prospective cohort of returning travelers.

Table S2. Staining panel for flow cytometry

Fig. S1. Flow cytometry manual gating strategy – T cell and B cell panel.

Fig. S2. MEFISTO model and cohort internal feature evaluation.

Fig. S3. Cohort external evaluation - comparison of plasma protein and PBMC profiles at Acute and Y1 compared to healthy controls.

Fig. S4. Comparison of MEFISTO Factor1 values for exposure groups on time points.

Fig. S5. Cumulative Response Score for malaria antigen specific IgG subclass response of Yman et al, BMC Medicine, 2019 data set.

Fig. S6. Leukocyte count adjusted cell counts.

Fig. S7. Directed Acyclic Graphs – DAGs and linear regression

### Materials and Methods

Directed Acyclic Graphs and linear regression models

To further investigate the causal relationship of our correlates, we use Directed Acyclic Graphs (DAG), also known as causal diagrams, to draw a graphic model based on the hypothesis and information we have. Based on the relationships between all the nodes in a DAG, one can find the minimal sufficient adjustment set for estimating the total effect of some exposure on some outcome. In a DAG, these adjustments set variables are enough to adjust for to close all backdoors relationship paths which is needed to identify the relationship between exposure and outcome.
Linear regression modelling was used to assess if correlated immune features can resemble the differences we observe when looking at previous exposure only.
For that, a simple model was fitted, were the immune response is explained by the exposure group as explanatory variable - reference model (1). The reference model was then compared with a model associated immune features as explanatory variable without adjustment (2) and with the minimal adjustment set determined in the corresponding DAG (3).
Figure S7A shows the DAG that was used to determine if iRBCs drive acute IFN-gamma levels with the minimal adjustment set of intermediate monocytes and IgG3 response (3).
Comparing the R2 value of the reference model (1) to the adjusted immune parameter model (3) gives insight if the model explains the data in better (greater R2 value) or worse (lower R2 value).

### Table S1. Characteristics of the Malaria Travelers Cohort

| | Primary infected (N=17) | Previously exposed (N=36) | All (N=53) | p value |
|---|---|---|---|---|
| **Gender** | | | | 0.551[1] |
| female | 4 (23.5%) | 6 (16.7%) | 10 (18.9%) | |
| male | 13 (76.5%) | 30 (83.3%) | 43 (81.1%) | |
| **Age [years]** | | | | 0.370[2] |
| Mean (SD) | 37.00 (10.68) | 40.11 (10.73) | 39.11 (10.71) | |
| Median (Min, Max) | 34 (20, 60) | 39 (27, 69) | 38 (20, 69) | |
| **CMV serology status** | | | | 0.008[1] |
| negative | 5 (29.4%) | 1 (3.1%) | 6 (12.2%) | |
| positive | 12 (70.6%) | 31 (96.9%) | 43 (87.8%) | |
| **Cumulative time of residency in malaria endemic area [years]** | | | | < 0.001[2] |
| Mean (SD) | 0.59 (1.12) | 25.53 (7.02) | 17.48 (13.16) | |
| Median (Min, Max) | 0 (0, 3) | 25 (14, 39) | 20 (0, 39) | |
| **Time since permanent residency in malaria endemic area [years]** | | | | |
| Mean (SD) | - | 13.39 (11.32) | 13.39 (11.32) | |
| Median (Min, Max) | - | 11.5 (0, 46) | 11.5 (0, 46) | |
| **Self-reported symptom onset before acute sample [days]** | | | | 0.954[2] |
| Mean (SD) | 4.71 (2.93) | 4.94 (3.47) | 4.87 (3.28) | |
| Median (Min, Max) | 4 (0, 12) | 5 (1,21) | 5 (, 21) | |
| Missing | 1 | 1 | 2 | |
| **Fever at admission (≥ 38°C)** | | | | 0.279[1] |
| no | 1 (5.9%) | 6 (16.7%) | 7 (13.2%) | |
| yes | 16 (94.1%) | 30 (83.3%) | 46 (86.8%) | |
| **Body temperature at admission [°C]** | | | | 0.062[2] |
| Mean (SD) | 38.86 (1.29) | 38.21 (1.23) | 38.42 (1.27) | |
| Median (Min, Max) | 39.0 (36.1, 40.5) | 38.0 (36.2, 40.6) | 38.4 (36.1, 40.6) | |
| **Leukocytes at admission ($\times 10^9$/L) (normal range 3.8–8.8)** | | | | 0.086[2] |
| Mean (SD) | 4.46 (1.60) | 6.26 (5.43) | 5.66 (4.58) | |
| Median (Min, Max) | 4.2 (2.1, 7.5) | 5.25 (1.9, 35.0) | 5.0 (1.9, 35.0) | |
| Missing | 0 | 2 | 2 | |
| **C-reactive protein at admission (mg/L) (normal range 0–3)** | | | | 0.441[2] |
| Mean (SD) | 111.53 (58.00) | 106.69 (83.72) | 108.27 (75.72) | |

| | Primary infected (N=17) | Previously exposed (N=36) | All (N=53) | p value |
|---|---|---|---|---|
| Median (Min, Max) | 105 (29, 237) | 88 (6, 294) | 90 (6, 294) | |
| Missing | 0 | 1 | 1 | |
| **Duration of hospital admission [days]** | | | | 0.002[2] |
| Mean (SD) | 4.76 (4.96) | 3.28 (5.26) | 3.75 (5.16) | |
| Median (Min, Max) | 3 (1, 23) | 2 (0, 32) | 3 (0, 32) | |
| **ICU admission** | | | | 0.962[1] |
| no | 16 (94.1%) | 34 (94.4%) | 50 (94.3%) | |
| yes | 1 (5.9%) | 2 (5.6%) | 3 (5.7%) | |
| **Severe malaria with at least one of WHO criteria (2010) or hyper-parasitemia >2%** | | | | 0.034[1] |
| no | 8 (53.3%) | 28 (82.4%) | 36 (73.5%) | |
| yes | 7 (46.7%) | 6 (17.6%) | 13 (26.5%) | |
| Missing | 2 | 2 | 4 | |
| **Severe malaria according to WHO 2014 without hyper-parasitemia as a single criterion (60)** | | | | 0.543[1] |
| no | 16 (94.1%) | 32 (88.9%) | 48 (90.6%) | |
| yes | 1 (5.9%) | 4 (11.1%) | 5 (9.4%) | |

1. Pearson's Chi-squared test
2. Kruskal-Wallis rank sum test

57

### Table S2. Staining panel for flow cytometry

| Antibody specificity | Fluorochrome conjugate | Antibody clone | Source |
|---|---|---|---|
| **Landscape panel** | | | |
| CD14 | BB700 | MϕP9 | BD |
| CD57 | BB515 | NK-1 | BD |
| γδTCR | PECy7 | 11F2 | BD |
| CD38 | PECy5 | HIT2 | BD |
| CD25 | PE-CF594 | M-A251 | BD |
| Vδ2 | PE | b6 | BD |
| CD3 | APC-H7 | SK7 | BD |
| CD16 | AF700 | 3G8 | BD |
| CCR7 | AF647 | 3D12 | BD |
| CD56 | BV786 | NCAM16.2 | BD |
| CD19 | BV711 | SJ25C1 | BD |
| CD45RA | BV650 | HI100 | BD |
| HLA-DR | BV605 | G46-6 | BD |
| Aqua LIVE/DEAD | BV510 channel | | ThermoFisher |
| CD127 | BV421 | HIL-7R-M21 | BD |
| CD8 | BUV737 | SK1 | BD |
| CD4 | BUV395 | SK3 | BD |
| **Restimulation experiment** | | | |
| CD107a (LAPM-1) | FITC | H4A3 | Biolegend |
| CD3 | PE-CF594 | UCHT1 | BD |
| γδ TCR | PE-Cy7 | 11F2 | BD |
| Vδ2 | PE | B6 | Biolegend |
| IFNγ | APC-Cy7 | B27 | Biolegend |
| IL10 | APC | JES3-19F1 | Biolegend |
| IL17 | BV711 | BL168 | Biolegend |
| TNF | BV605 | Mab11 | Biolegend |
| Aqua LIVE/DEAD | BV510 channel | | ThermoFisher |

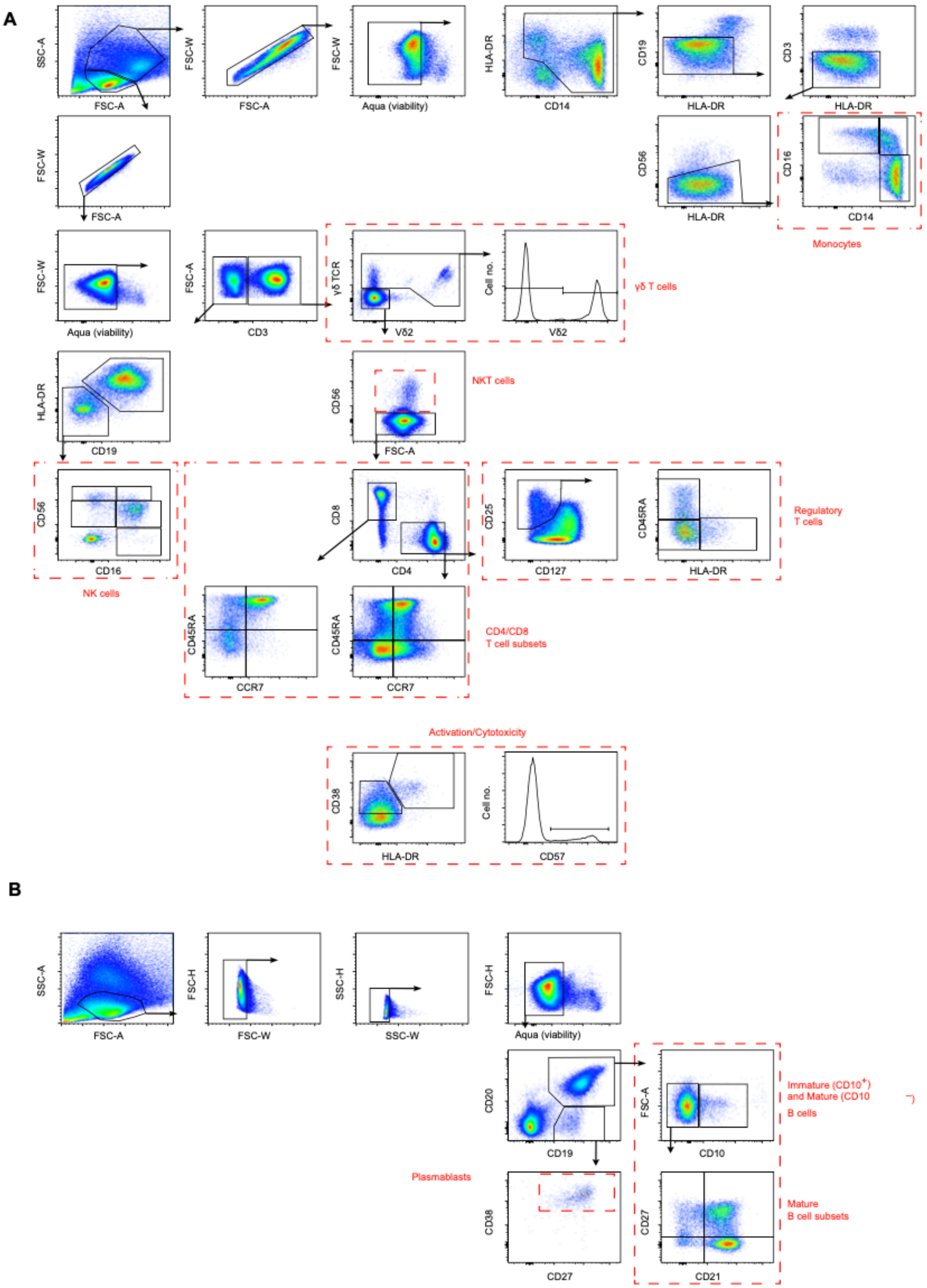


**Fig. S1. Flow cytometry manual gating strategy – T cell and B cell panel.** (**A**) 17-marker panel with manual gating of cell populations included in the systems analysis. The included populations are indicated by dashed red boxes. **(B)** 13-marker panel focused on

B cells. Manual gating of populations included in the analysis indicated by dashed red boxes. The B cell data was generated in Sundling et al., JCI insight, 2019.

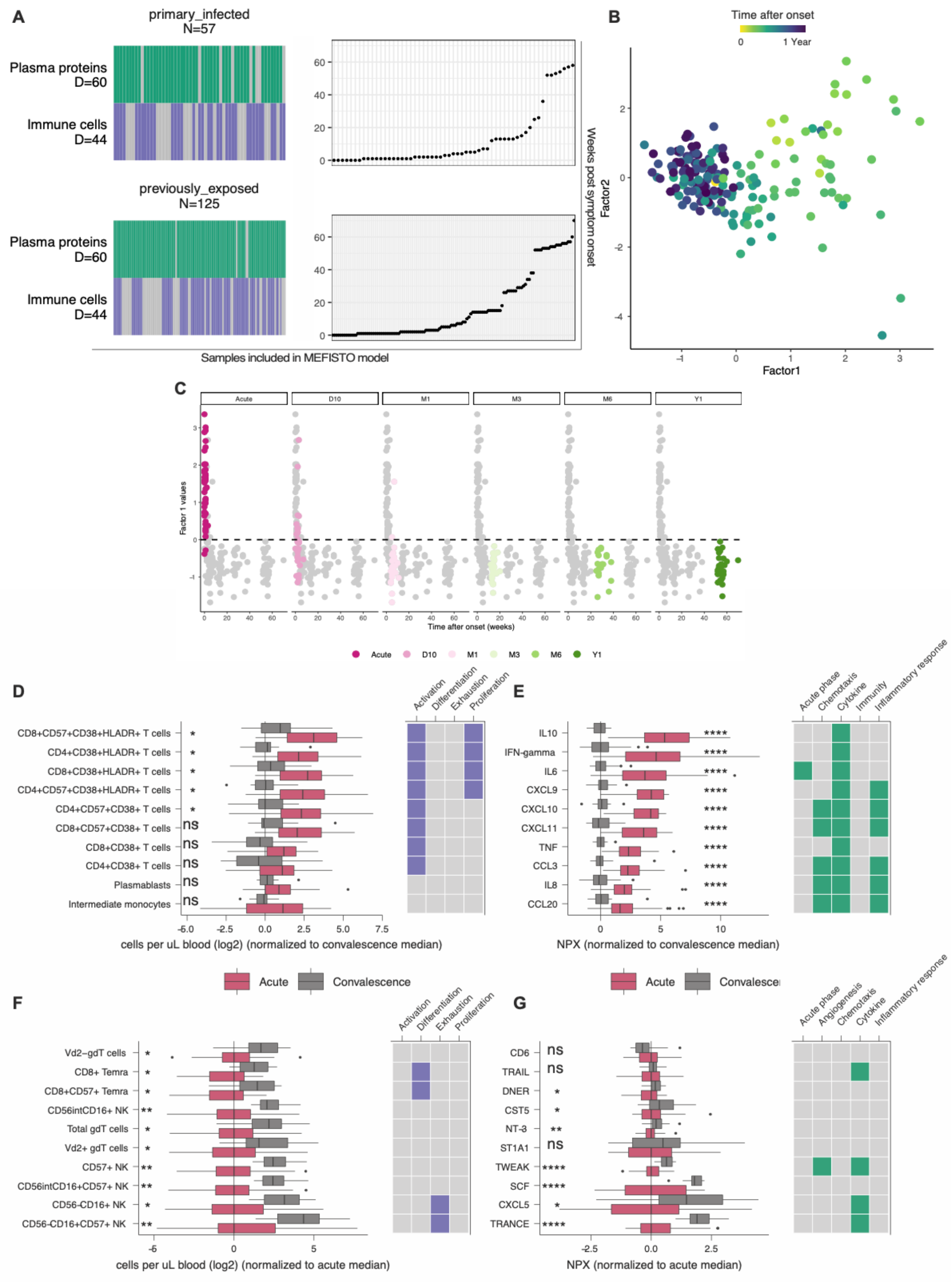
A
primary_infected
N=57
Plasma proteins
D=60
Immune cells
D=44
previously_exposed
N=125
Plasma proteins
D=60
Immune cells
D=44
Weeks post symptom onset
Samples included in MEFISTO model
B
Time after onset
0
1 Year
Factor2
Factor1
C
Acute
D10
M1
M3
M6
Y1
Factor 1 values
Time after onset (weeks)
D
CD8+CD57+CD38+HLADR+ T cells
CD4+CD38+HLADR+ T cells
CD8+CD38+HLADR+ T cells
CD4+CD57+CD38+HLADR+ T cells
CD4+CD57+CD38+ T cells
CD8+CD57+CD38+ T cells
CD8+CD38+ T cells
CD4+CD38+ T cells
Plasmablasts
Intermediate monocytes
Activation
Differentiation
Exhaustion
Proliferation
cells per uL blood (log2) (normalized to convalescence median)
E
IL10
IFN-gamma
IL6
CXCL9
CXCL10
CXCL11
TNF
CCL3
IL8
CCL20
Acute phase
Chemotaxis
Cytokine
Immunity
Inflammatory response
NPX (normalized to convalescence median)
Acute
Convalescence
F
Vd2−gdT cells
CD8+ Temra
CD8+CD57+ Temra
CD56intCD16+ NK
Total gdT cells
Vd2+ gdT cells
CD57+ NK
CD56intCD16+CD57+ NK
CD56-CD16+ NK
CD56-CD16+CD57+ NK
cells per uL blood (log2) (normalized to acute median)
G
CD6
TRAIL
DNER
CST5
NT-3
ST1A1
TWEAK
SCF
CXCL5
TRANCE
Acute phase
Angiogenesis
Chemotaxis
Cytokine
Inflammatory response
NPX (normalized to acute median)

**Fig. S2. MEFISTO model and cohort internal feature evaluation.** (**A**) Overview of available data modality (colored) and missing data (grey) data in each patient sample used as input for MEFISTO. (**B**) Latent space of Factor1 and Factor2, colored by time after onset up to one year. (**C**) Factor1 values for each sample time point. Internal comparison of Top10 positive Factor1 associated features for (**D**) cell subsets and (**E**) plasma proteins at acute (red) time point with convalescence (grey) at 6-12month. Feature annotation based on functionality marker surface expression on immune cells (purple) and UniProt keywords associated with proteins (green). Similarly, internal comparison of Top10 negative Factor1 associated features for cell subsets (**F**) and plasma proteins (**G**) at convalescence compared to acute time point. Box plots visualize values, cohort internally normalized to acute/convalescence median of immune cells and plasma proteins, respectively. Statistically differences were assessed using the non-parametric Wilcoxon-test with correction for multiple testing, adjusted p values (FDR < 0.05). *p < 0.05, **p < 0.01, ***p < 0.001

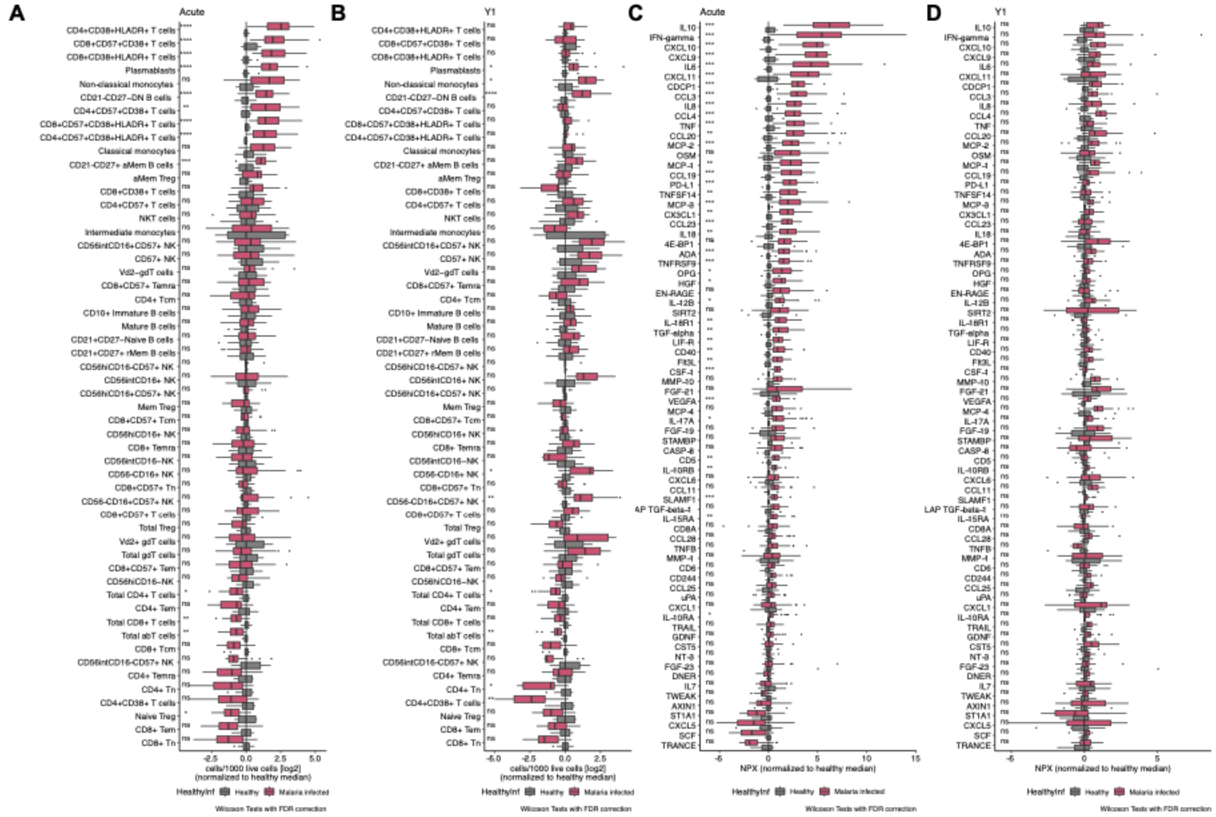

**Fig. S3. Cohort external evaluation - comparison of plasma protein and PBMC profiles at Acute and Y1 compared to healthy controls.** Immune cell subset counts (cells/1000 live cells) at **(A)** acute and (**B**) convalescence (6-12 month after disease) of malaria patients (red) compared to healthy controls (grey). Relative plasma protein levels (NPX) at **(C)** acute and (**D**) convalescence (6-12 month after disease) compared to healthy controls. Box plots visualize values, normalized to the healthy control median of immune cells and plasma proteins, respectively. Statistical differences were assessed using the non-parametric Wilcoxon-test with FDR correction for multiple testing. *p < 0.05, **p < 0.01, ***p < 0.001

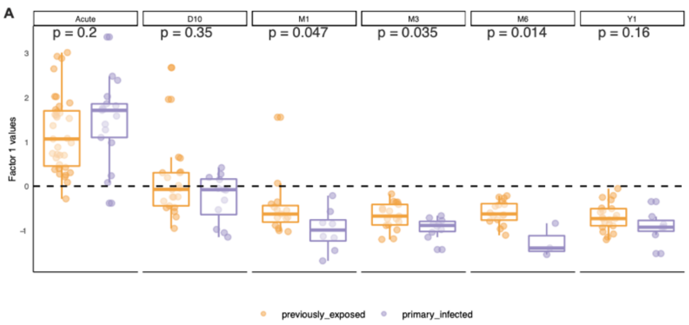


**Fig. S4. Comparison of MEFISTO Factor1 values for exposure groups on time points.** (A) Comparison of Factor1 values magnitudes between the two groups of primary infected (purple) and previously exposed (orange) individuals. Statistical differences were assessed using the non-parametric Wilcoxon-test. *p < 0.05, **p < 0.01, ***p < 0.001

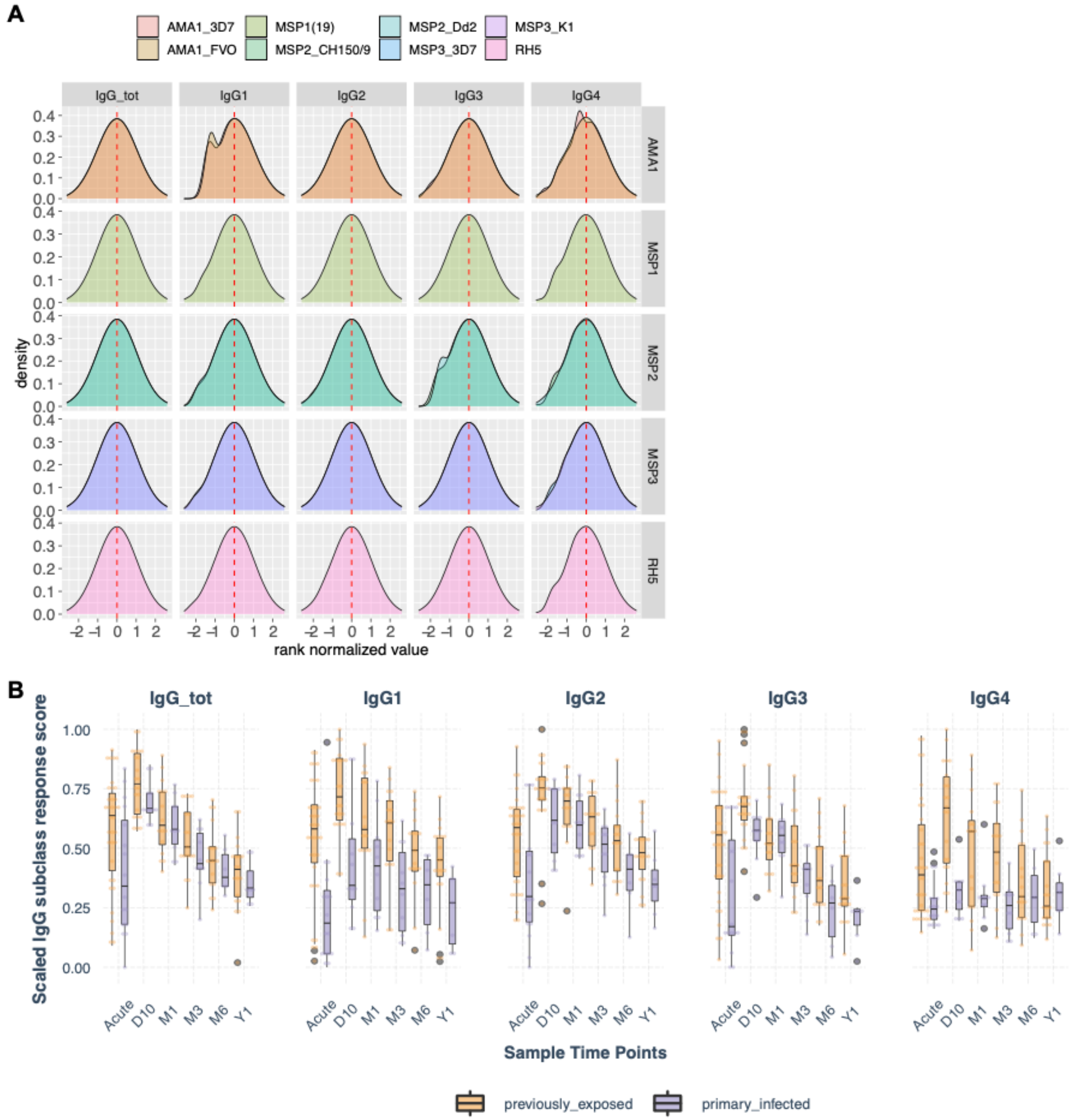


**Fig. S5. Cumulative malaria antigen specific IgG subclass response of Yman et al 2019 data set.** (**A**) Density plot of rank normalized data from repurposed data set, showing normally distributed data for total IgG, each IgG subclass (columns) and five parasite-antigen (rows) and the specific antigen (colors). (**B**) Scaled IgG subclass response score (denoted in manuscript as CRS) for each IgG subclass over time, presenting score of subclass-breadth response.

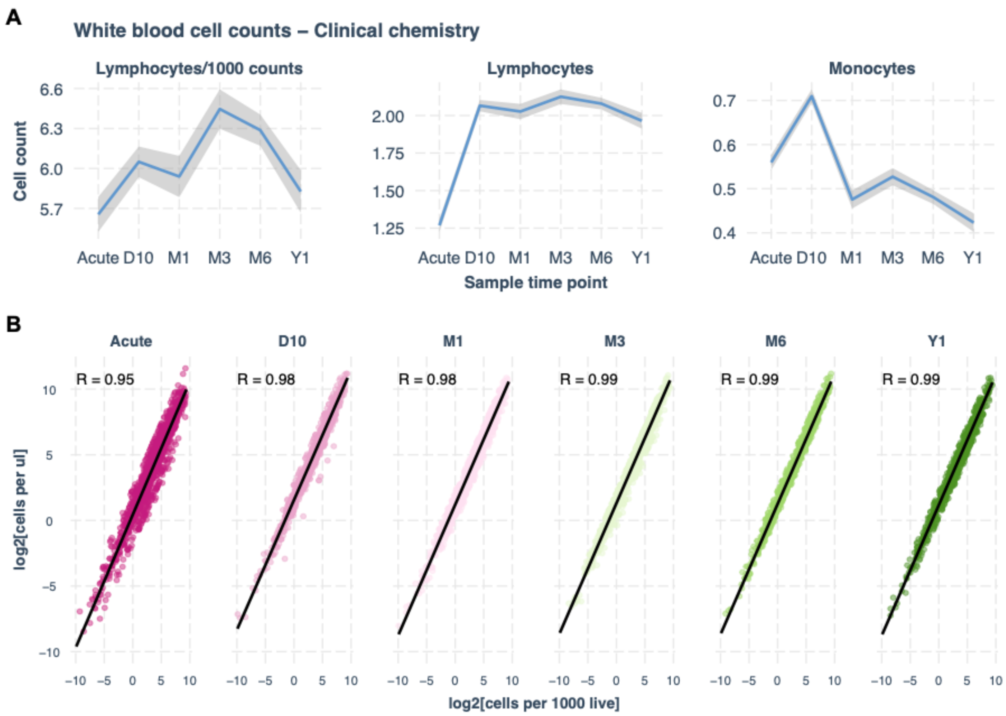


**Fig. S6. Leukocyte count adjusted cell counts.** (**A**) Clinical Chemistry white blood cell counts in cohort samples over time. Left graph indicates cell counts normalized to total number of live cells in sample. Middle and right graphs indicate counts based on cells per µl whole blood as measured in the hospital clinical diagnostics lab. (**B**) Correlation of white blood cells normalized per µl blood data with flow cytometry data normalized to cells per 1000 live cells.

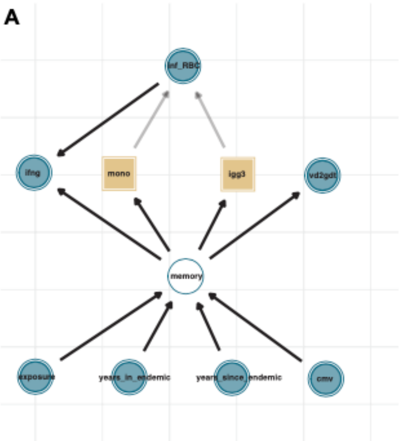


| | (1) PrevExposure | (2) iRBC | (3) iRBC \| IgG3 + int. monocytes |
|---|---|---|---|
| (Intercept) | 12.006*** | 9.731*** | 14.913*** |
| | (0.955) | (0.946) | (1.603) |
| group | −3.801** | | |
| | (1.257) | | |
| inf_rbc | | 0.043 | −0.173 |
| | | (0.316) | (0.265) |
| IgG3 | | | −5.610* |
| | | | (2.185) |
| mono_int | | | −0.642* |
| | | | (0.273) |
| Num.Obs. | 26 | 26 | 26 |
| R2 | 0.276 | 0.001 | 0.386 |
| R2Adj. | 0.246 | −0.041 | 0.302 |
| AIC | 137.7 | 146.0 | 137.4 |
| BIC | 141.4 | 149.8 | 143.7 |
| Log.Lik. | −65.827 | −70.013 | −63.687 |
| F | 9.141 | 0.019 | 4.606 |

+p<0.1,*p<0.05,**p<0.01,***p<0.001

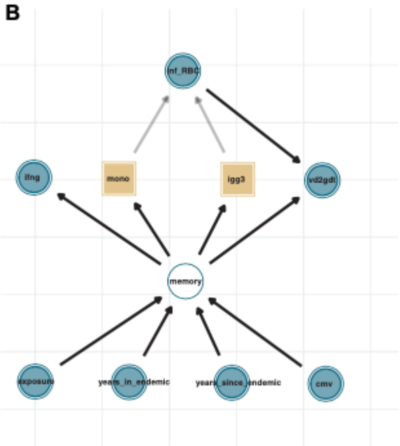


| | (1) PrevExposure | (2) iRBC | (3) iRBC \| IgG3 + int. monocytes |
|---|---|---|---|
| (Intercept) | 3.932** | 1.854+ | 5.043 |
| | (0.809) | (0.955) | (3.748) |
| group | −2.615* | | |
| | (1.085) | | |
| inf_rbc | | 0.298 | −0.084 |
| | | (0.316) | (0.570) |
| IgG3 | | | −2.114 |
| | | | (2.745) |
| mono_int | | | −0.321 |
| | | | (0.615) |
| Num.Obs. | 9 | 9 | 9 |
| R2 | 0.453 | 0.113 | 0.285 |
| R2Adj. | 0.375 | −0.014 | −0.145 |
| AIC | 37.9 | 42.3 | 44.4 |
| BIC | 38.5 | 42.9 | 45.3 |
| Log.Lik. | −15.971 | −18.149 | −17.180 |
| F | 5.802 | 0.891 | 0.663 |

+p<0.1,*p<0.05,**p<0.01,***p<0.001

| | (1) PrevExposure | (2) iRBC | (3) iRBC \| IgG1 + int. monocytes |
|---|---|---|---|
| (Intercept) | 3.932** | 1.854+ | 6.174+ |
| | (0.809) | (0.955) | (2.503) |
| group | −2.615* | | |
| | (1.085) | | |
| inf_rbc | | 0.298 | −0.323 |
| | | (0.316) | (0.386) |
| IgG1 | | | −6.066* |
| | | | (2.157) |
| mono_int | | | −0.135 |
| | | | (0.404) |
| Num.Obs. | 9 | 9 | 9 |
| R2 | 0.453 | 0.113 | 0.690 |
| R2Adj. | 0.375 | −0.014 | 0.504 |
| AIC | 37.9 | 42.3 | 36.8 |
| BIC | 38.5 | 42.9 | 37.8 |
| Log.Lik. | −15.971 | −18.149 | −13.417 |
| F | 5.802 | 0.891 | 3.710 |

+p<0.1,*p<0.05,**p<0.01,***p<0.001

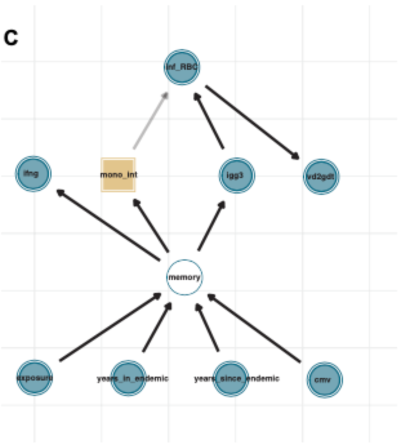


| | (1) PrevExposure | (2) IgG3 | (3) IgG3 \|int. monocytes |
|---|---|---|---|
| (Intercept) | 7.404*** | 8.468*** | 7.008*** |
| | (1.022) | (0.784) | (1.150) |
| group | −2.171+ | | |
| | (1.190) | | |
| IgG3 | | −5.903*** | −4.193+ |
| | | (1.478) | (1.814) |
| mono_int | | | 0.131 |
| | | | (0.266) |
| Num.Obs. | 19 | 19 | 9 |
| R2 | 0.164 | 0.484 | 0.498 |
| R2Adj. | 0.114 | 0.454 | 0.331 |
| AIC | 89.2 | 80.0 | 36.6 |
| BIC | 92.0 | 82.9 | 37.4 |
| Log.Lik. | −41.604 | −37.014 | −14.286 |
| F | 3.327 | 15.955 | 2.982 |

+p<0.1,*p<0.05,**p<0.01,***p<0.001

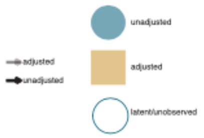


**Fig. S7. Directed Acyclic Graphs and linear regression.** Directed acyclic graphs based on results and assumptions (graphic) with minimal sufficient adjustment set (sand color) and linear regression model summary of (1) reference model, (2) un-adjusted model and (3) DAG-based adjusted model. (**A**) DAG with minimal sufficient adjustment set for estimating the total effect of parasitemia (inf_rbc) on acute IFN-gamma levels, adjusted for parasite-specific IgG3 and intermediate monocytes. (**B**) DAG with minimal sufficient adjustment set for estimating the total effect of parasitemia (inf_rbc) on $V\delta2^+$ $\gamma\delta$ T cell numbers at the expansion time point D10 (vd2gdt), adjusted for parasite-specific IgG3 and intermediate monocytes and in a separate model with parasite-specific IgG1 replacing IgG3. (**C**) DAG with minimal sufficient adjustment set for estimating the total effect of parasite specific IgG3 on $V\delta2^+$ $\gamma\delta$ T cell numbers at the expansion time point D10 (vd2gdt), adjusted for intermediate monocyte levels. Model statistics assessed with ANOVA; $+p < 0.1$, $*p < 0.05$, $**p < 0.01$, $***p < 0.001$
